# Spatial scale of indentation explains shift in ratio between spinal cord gray and white matter stiffness

**DOI:** 10.64898/2026.04.20.719143

**Authors:** Oskar Neumann, Harsh Vardhan Surana, Maik Hintze, Stefanie Kuerten, Thomas Franz, Rahul Gopalan Ramachandran, Paul Steinmann, Silvia Budday

**Affiliations:** Institute of Continuum Mechanics and Biomechanics, Friedrich-Alexander-Universität Erlangen-Nürnberg, Dr.-Mack-Straße 81, Fürth, 90762, Germany; Institute of Neuroanatomy, Faculty of Medicine, University of Bonn and University Hospital Bonn, Nußallee 10, Bonn, 53115, Germany; Institute of Anatomy and Cell Biology, Friedrich-Alexander Universität Erlangen-Nürnberg, Krankenhausstr. 9, Erlangen, 91054, Germany; Biomedical Engineering Research Centre, Division of Biomedical Engineering, Department of Human Biology, Faculty of Health Sciences, University of Cape Town, Observatory, Cape Town, 7925, South Africa; Bioengineering Science Research Group, Department of Mechanical Engineering, Faculty of Engineering and Physical Sciences, University of Southampton, Highfield, Southampton, SO17 1BJ, United Kingdom; Institute of Applied Mechanics, Friedrich-Alexander-Universität Erlangen-Nürnberg, Egerlandstraße 5, Erlangen, 91058, Germany

**Keywords:** Tissue mechanics, Spinal cord, gray matter, White matter, Indentation, Multiscale testing, Mechanical properties

## Abstract

The structural integrity of spinal cord tissue and the transmission of mechanical stimuli across the different levels of tissue microarchitecture and varying spatial scales of mechanical loading challenge experimental and computational efforts to accurately model, simulate and interpret tissue mechanics, leading to conflicting findings in existing literature. Here, we demonstrate that the bead size used in spherical indentation tests significantly affects the stiffness ratio of spinal cord gray to white matter, a dependence which we only observe on the transverse plane and not the coronal plane of the tissue. Our study reveals a shift in stiffness ratio such that for smaller spherical indenters gray matter is stiffer than white matter, while for larger indenters, white matter is stiffer than gray matter. The mean relative change from the 100 *µ*m bead to the 500 *µ*m bead differed between anatomical planes, with transverse sections showing a decrease in gray matter (−13.3%) and an increase in white matter stiffness (+26.9%), accompanied by a reduction in the gray-to-white matter stiffness ratio from 1.07 to 0.76, whereas coronal sections exhibited increases in both gray (+21.0%) and white matter (+33.8%), along with a change in the ratio from 0.99 to 1.14. These findings contribute to explaining previously contradictory results in the literature and underscore the relevance of spatial scales in mechanical characterization studies.

## 1. Introduction

Scientific investigations into the mechanical properties of spinal cord tissue have expanded significantly over recent decades. Despite valuable advances in the mechanical characterization of central nervous system (CNS) tissue in the spinal cord, there are large differences and perceived inconsistencies in the reported tissue stiffnesses, especially when it comes to indentation tests - Atomic Force Microscopy (AFM), nanoindentation, or comparable methods. These inconsistencies can be attributed to variations in sample preparation, experimental conditions, methods for data post-processing, and testing procedures [1, 2, 3]. A frequently addressed research question in this context is which of the CNS tissue types – gray or white matter tissue – is more compliant when measured by indentation [4]. Some studies report stiffer white matter tissue [5, 6, 7, 8, 9, 10], others observed the opposite [11, 12, 13, 14, 15, 16, 17, 18], still others see no difference in the mechanical response between white and gray matter tissue [19, 20]. Qian et al. emphasizes that the mechanical heterogeneity of CNS tissue is related to its biological heterogeneity [21]. A closer examination of the available literature reveals that although these studies all report on the similar mechanical loading scenario, indentation tests, they differ considerably in experimental parameters. Previous studies have demonstrated that the measured mechanical properties of white and gray matter can vary depending on factors such as loading rate, maximum applied force, and *post mortem* time [4, 22], as well as sample preparation [22] and the experimental model employed (*in vitro, in situ, in vivo*, etc.) [23]. While acknowledging this multitude of potentially influencing experimental parameters, the present study focuses specifically on the size of the applied spherical indenter in order to investigate its influence on the measured mechanical properties of spinal cord white and gray matter tissue as well as their ratio. In contrast, other biological materials such as bone and cartilage, which are relatively more structurally uniform at the spatial scales typically probed by indentation compared with CNS tissue, were found to have similar mechanical properties at different spatial scales [24, 25]. Interestingly, Wahlsten et al. found no significant differences in the stiffness of human skin tissue measured by microindentation (spherical bead with a diameter of 200 *µ*m) or AFM (spherical bead with a diameter of 6.10 *µ*m) [26]. These findings suggest at least a correlative, if not a causal, relationship between the contradictory results reported in spinal cord tissue indentation tests and the varying indenter sizes used. In contrast to more homogeneous tissues, such as bone, cartilage, and skin, which exhibit consistent mechanical properties across spatial scales, highly heterogeneous tissues like brain and spinal cord display considerable variability across the different studies.

An elaborate understanding of how the mechanical response of CNS tissue is controlled from cellular scales (tens of micrometers) through tissue scales (hundreds of micrometers) to organ scales (centimeter range) could enhance scientific endeavors in simulating natural and enforced changes in the mechanical state of CNS tissue [26], which have been shown to be linked with various pathologies [27]. In recent years, substantial progress has been made in developing computational models to simulate mechanobiological processes in spinal cord and brain tissue across spatial scales, ranging from molecules to the tissue or even organ scale [28]. For instance, Biglari et al. investigated the deformation of the vertebral body, intervertebral disc and spinal cord tissue during injuries of the thoracic spinal cord using patient-specific finite element models [29]. In the context of such computational models, shifts in the ratio between gray and white matter stiffness can vastly impact the simulation outcome [30], which highlights the importance of understanding corresponding discrepancies in the literature. To isolate one of the multiple factors influencing the measured mechanical properties of spinal cord tissue, here, we focus exclusively on one mechanical testing modality – spherical indentation. First, we review and analyze experimental results in the literature regarding indentersize-dependent mechanical properties, with special focus on the ratio of gray to white matter stiffness. To experimentally investigate the influence of the indenter size on this ratio, we then developed a reliable, consistent and repeatable protocol to test porcine spinal cord tissue with spherical indenters of different sizes. In the course of this study, we performed 2861 indentations on 32 samples of 32 different porcine spinal cords.

## 2. Results

### 2.1. Correlation analysis of literature data

In a first step, we assessed possible influences of different test conditions on stiffness ratios reported in the literature. Specifically, we tested possible correlations between the ratio of gray to white matter stiffness and four different experimental parameters: the diameter of the indenter, the maximum force, the indentation depth, and the loading rate. The corresponding data of comparable studies are summarized in Tab. S1.

Fig. S1 shows the results of a linear regression analysis for the four different experimental parameters. The corresponding results are summarized in Tab. 1. Only the correlation between indenter size and stiffness ratio shows statistical significance (*P* = 0.032). This observation motivated us to study the effect of the indenter size on stiffness values measured in gray and white matter spinal cord tissue and their ratio in more detail, as described in the following.

**Table 1:** Results for regression analyses shown in Fig. S1.

| Correlation test | $R^2$ | P |
| --- | --- | --- |
| indenter diameter vs. ratio | 0.2203 | 0.0318 |
| max force vs. ratio | 0.2988 | 0.1021 |
| indentation depth vs. ratio | 0.0822 | 0.3663 |
| loading rate vs. ratio | 0.1162 | 0.1806 |

$R^2$ , coefficient of determination; P, two-tailed Wald $t$ -test for zero slope.

### 2.2. Variability of mechanical properties between samples

Fig. 1a illustrates the sample-wise mean apparent moduli of tests with increasing indenter diameter (D) from D = 20 (blue), to 100 (orange), and 500 (gray) *µ*m on transverse sections (Protocol A) of spinal cord gray (dark) and white (light) matter. The results show mean values of 1.57 ± 1.23, 1.15 ± 0.43, and 1.14 ± 0.77 kPa for indentations on spinal cord gray and 1.73 ± 1.18, 1.16 ± 0.49, and 1.76 ± 1.14 kPa for indentations on spinal cord white matter tissue using the smallest, intermediate and largest indenter diameters, respectively. Overall, inter-sample variability was substantial: mean stiffness spanned from 0.99 (Sample 3) to 4.08 kPa (Sample 2) and from 0.74 (Sample 3) to 3.81 kPa (Sample 2) for the smallest indenter, from 0.69 (Sample 3) to 1.85 kPa (Sample 6) and from 0.69 (Sample 5) to 1.69 kPa (Sample 4) for the intermediate, and from 0.47 (Sample 3) to 2.28 kPa (Sample 7) and from 0.66 (Sample 2) to 3.61 kPa (Sample 6) for the largest indenter in spinal cord gray and white matter tissue, respectively.

**Figure 1:**
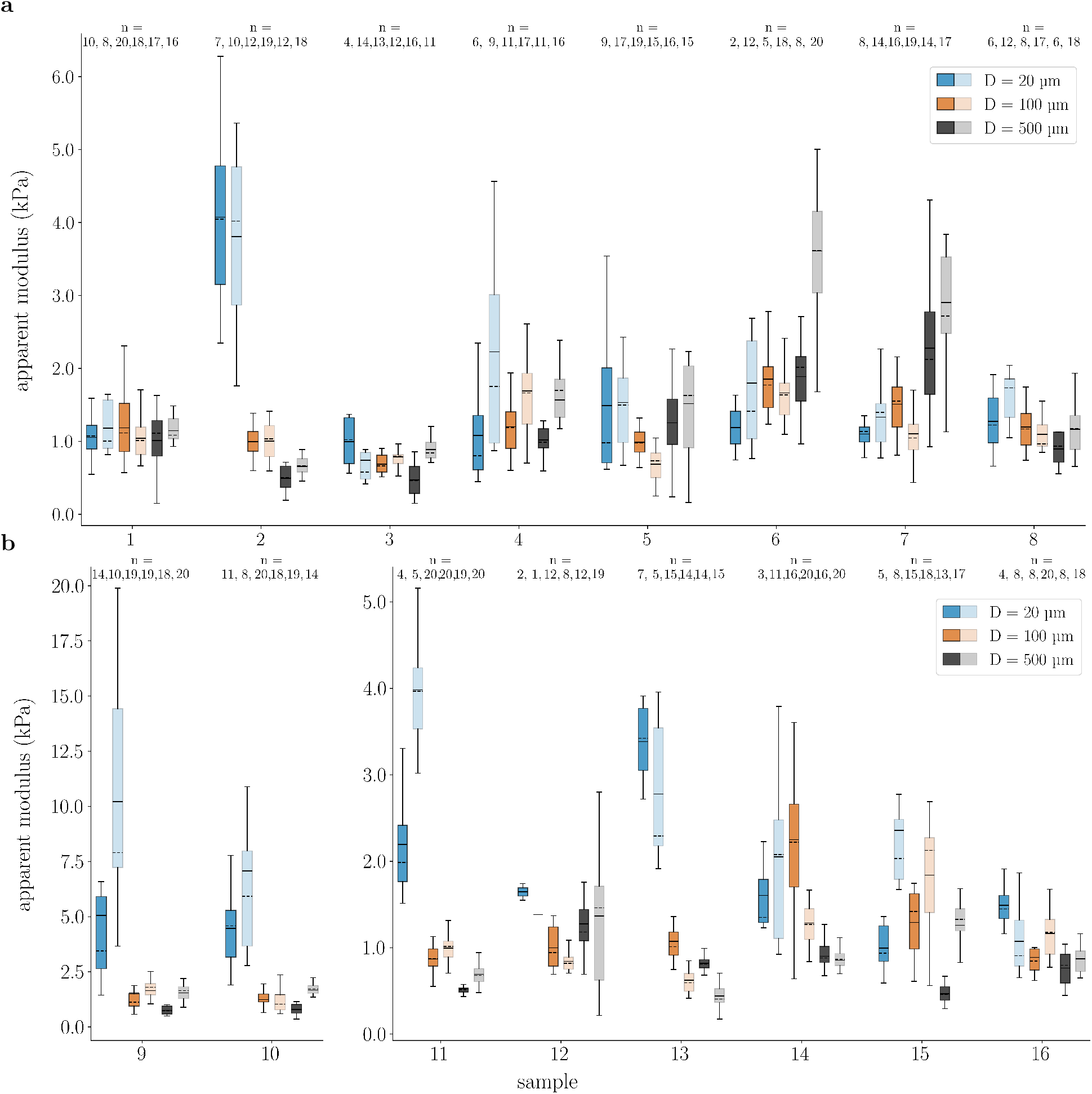
Apparent moduli for indentations across samples for three different indenter sizes. (**a**) Results for tests according to Protocol A, order of measurements: D = 20, D = 100, D = 500 *µ*m), transverse plane. (**b**) Results for tests according Protocol B, order of measurements: D = 500, D = 100, D = 20 *µ*m, transverse plane. Results are collectively clustered for each sample. Indentations with D = 20, 100 and 500 *µ*m are shown in blue, orange and gray, respectively. Lighter colors depict results for spinal cord white matter tissue, darker colors for spinal cord gray matter tissue. Boxes show the 25th (Q1), 50th (median) and 75th (Q3) percentiles with inter-quartile range (IQR: from Q1 to Q3); whiskers reach the most extreme data points within ± 1.5 × IQR of Q1 and Q3. Outliers outside this range are not displayed but were included in the analysis. Mean values are indicated as continuous lines, median values as dashed lines. n indicates the number of technical replicates.

When comparing the results for gray and white matter of different samples, we observe a strong coupling: if one tissue type is stiffer or softer compared to other samples, the other tissue type of this sample will follow the same trend. Sample 2 shows the stiffest mechanical response for both gray and white matter tissue for tests with the smallest indenter. For the intermediate diameter, the two samples with the softest mechanical response in spinal cord gray matter tissue (Sample 3 and 5) also exhibit the softest white matter tissue response. These correlations become even more evident for the largest diameter, where the mean apparent moduli of spinal cord gray and white matter tissue show similar trends across most samples – except for Sample 6, which is stiffer than Sample 7 in the white matter, but not in the gray matter.

The sample-wise results from tests with decreasing indenter diameters on transverse sections (Protocol B) in Fig. 1b confirm the trends observed from Protocol A (increasing indenter diameters on transverse sections). We measured mean values of 3.43 ± 3.09, 1.30 ± 0.72, and 0.77 ± 0.30 kPa for indentations on spinal cord gray and 4.36 ± 4.35, 1.28 ± 0.65, and 1.08 ± 0.54 kPa for indentations on spinal cord white matter tissue using the smallest, intermediate, and largest indenter diameters, respectively. Similar to Fig. 1a, we can observe a high degree of inter-sample variability: mean stiffness spanned from 1.00 (Sample 15) to 5.06 kPa (Sample 9) and from 1.07 (Sample 16) to 1.02 kPa (Sample 9) for the smallest indenter, from 0.87 (Sample 11) to 2.25 kPa (Sample 14) and from 0.62 (Sample 13) to 1.84 kPa (Sample 15) for the intermediate, and from 0.47 (Sample 15) to 1.26 kPa (Sample 12) and from 0.44 (Sample 13) to 1.66 kPa (Sample 10) for the largest indenter in spinal cord gray and white matter tissue, respectively.

Again, we can observe a strong coupling between gray and white matter stiffness values. Sample 14 shows the highest overall stiffness for both spinal cord gray and white matter tissue for the largest diameter, while Sample 16 shows the stiffest response for both regions and the smallest diameter.

Similar results for Protocol C (refined analysis of larger indenters) and D (indentations on coronal sections) can be found in the Supplementary Material in Fig. S2a and Fig. S2a, respectively.

### 2.3. Indenter-size-dependent stiffness values and ratios of spinal cord gray to white matter tissue

We now assess the indenter-size dependence of spinal cord gray and white matter tissue stiffness values as well as their ratio – when averaging over all tested samples.

The results of tests with increasing indenter diameters on transverse sections (Protocol A) show a decrease in gray matter stiffness by a factor of 0.7 from the smallest to the intermediate diameter and a similar stiffness for the intermediate and the largest diameters (Fig. 2a and Tab. S3, first row of upper table). The normalized apparent moduli (see Eq. 3) yield an increase by 15% from the smallest to the intermediate and a decrease by 21% (*P* = 1.2 · 10^−5^) from the intermediate to the largest indenter diameter (Fig. 2e and Tab. 2, first row). On the contrary, we observe a decrease in spinal cord white matter tissue stiffness (Tab. S3, first row of lower table) by a factor of 0.7 (*P* = 0.0001) from the smallest to the intermediate diameter and an increase by a factor of 1.5 from the intermediate to the largest diameter (*P* = 9.6 · 10^−7^). This trend is also reflected in normalized tissue stiffness values (Fig. 2e and Tab. 2, first row) with a decrease by 8% from the smallest to the intermediate and an increase by 17% (*P* = 0.0001) from the intermediate to the largest indenter diameter.

**Figure 2:**
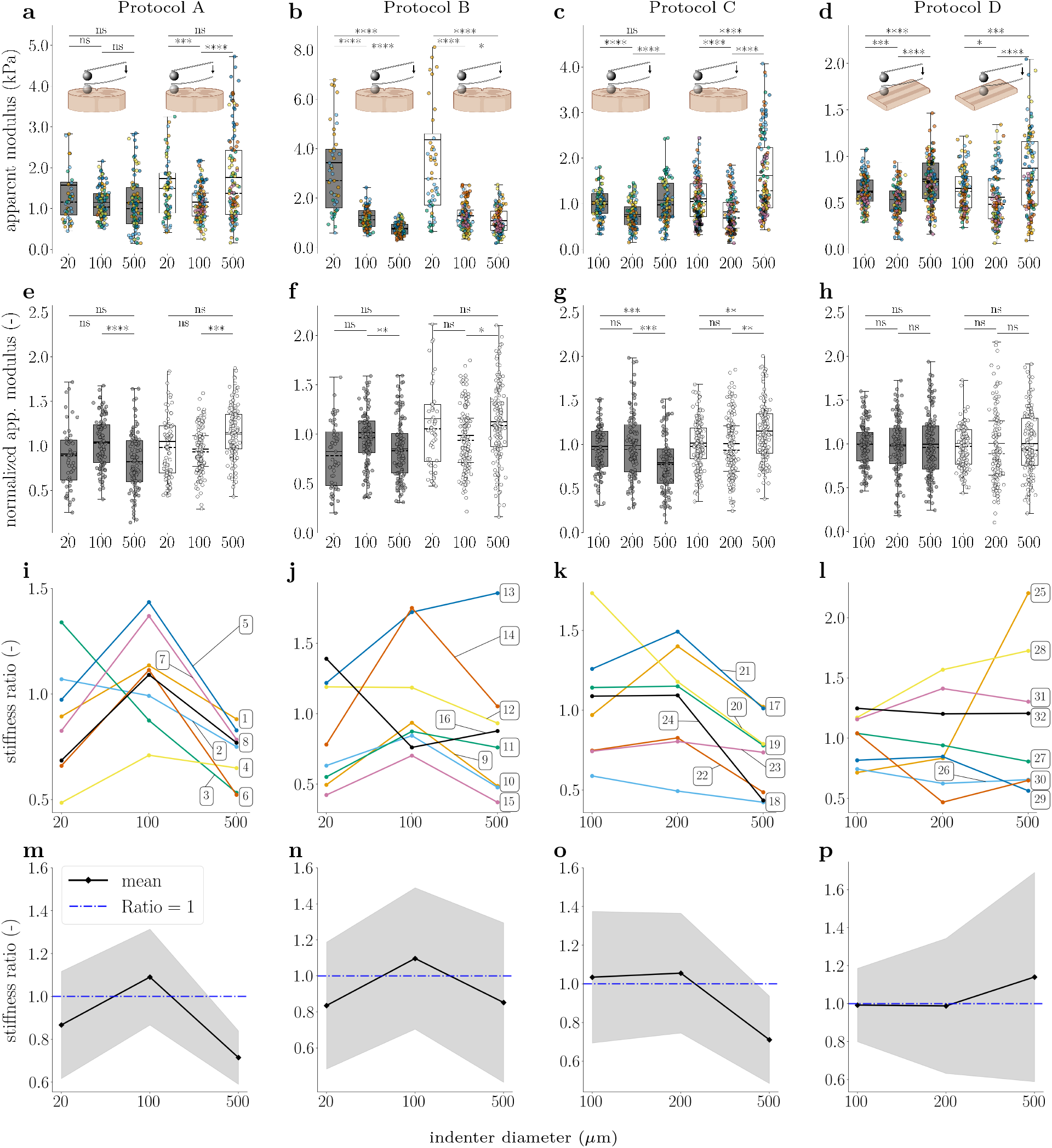
Indenter diameter influences ratio of gray to white matter stiffness. (**a, e, i, m**) results for Protocol A. (**b, f, j, n**) results for Protocol B. (**c, g, k, o**) results for Protocol C. (**d, h, l, p**) results for Protocol D. (**a**)-(**d**) Apparent moduli of technical replicates with color-coding. Each color represents one biological replicate. (**e**)-(**h**) normalized apparent moduli of technical replicates. gray and white boxes show the results of indentations on spinal cord gray and white matter tissue, respectively. Boxes show the 25th (Q1), 50th (median) and 75th (Q3) percentiles with inter-quartile range (IQR: from Q1 to Q3); whiskers reach the most extreme data points within ± 1.5 × IQR of Q1 and Q3. Outliers outside this range are not displayed but were included in the analysis. Data points are indicated as circles, mean values as continuous lines, and median values as dashed lines. The number of technical replicates (n) per boxplot is as follows: (**a, e**) n = 52, 104, 100, 96, 135, 131; (**b, f**) n = 50, 125, 119, 56, 137, 143; (**c, g**) n = 106, 110, 109, 157, 153, 147; (**d, h**) n = 135, 135, 156, 130, 139, 136. (**i**)-(**l**) Changes in ratio of spinal cord gray to white matter stiffness across different indenter diameters for individual biological replicates with sample numbers noted at the corresponding curves. Colors correspond to the scatter plot of apparent moduli of individual biological replicates in row one. (**m**)-(**p**) Mean ratio values of all biological replicates across different indenter diameters in black with standard deviation in gray. The blue dashed line indicates a ratio of one. The spinal cord inset sketches were created using BioRender.

**Table 2:** Changes in normalized mean apparent modulus across protocols.

| sequence of indenter diameter | anatomical plane | gray matter |  |  | white matter |  |  |
| --- | --- | --- | --- | --- | --- | --- | --- |
|  |  | 20 vs 100 | 100 vs 500 | 20 vs 500 | 20 vs 100 | 100 vs 500 | 20 vs 500 |
| PA: 20 → 100 → 500 | transverse | 15% | -21% | -9% | -8% | 17% | 8% |
| PB: 500 → 100 → 20 | transverse | 23% | -16% | 3% | -15% | 14% | -3% |
|  |  | 100 vs 200 | 200 vs 500 | 100 vs 500 | 100 vs 200 | 200 vs 500 | 100 vs 500 |
| PC: 100 → 200 → 500 | transverse | 1% | -20% | -19% | -1% | 14% | 14% |
| PD: 100 → 200 → 500 | coronal | 0% | 0% | 0% | 0% | 0% | 0% |

Colors indicate percentage differences as follows: turquoise, negative values; white, zero; magenta, positive values (shade intensity increases with absolute magnitude).

In Fig. 2i, every sample, except Samples 2 and 3, shows an increase in gray to white matter stiffness ratio from the smallest to the intermediate indenter diameter with an average increase in ratio of 0.39 (ranging from 0.24 to 0.54 for Samples 1, 4, 5, 6, 7 and 8). In contrast, the ratio decreases from the intermediate to the largest diameter for all samples with an average decrease of -0.38 (ranging from -0.60 to -0.10). Samples 1, 5, 6, 7 and 8 show an indenter-size dependent shift in the stiffness ratio with spinal cord white matter tissue being softer than gray matter for tests using the intermediate diameter and stiffer for the largest diameter. For Samples 2 and 3, this shift occurs already between the smallest and the intermediate diameter with a tendency of an even further decreasing ratio for the largest diameter. Sample 4 shows stiffer spinal cord white matter tissue for all indenter diameters, although the same general trend of a decreasing ratio from the intermediate to the largest diameter is observed, consistent with the other samples. The course of the overall mean ratios across all samples for each indenter diameter (Fig. 2m and Tab. 3, first row) reveals a shift from gray matter being stiffer than white matter (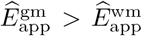, see Eq. 4) when using a 100 *µ*m diameter to white matter being stiffer than gray matter 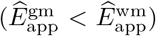 for tests using a 500 *µ*m diameter indenter. A fictitious diameter of 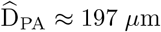 *µ*m can be estimated based on these data, at which the stiffness ratio of gray to white matter would be equal (ratio = 1). The corresponding slope of the connecting line is −9 · 10^−4^.

**Table 3:** Spinal cord gray to white matter tissue mean stiffness ratios across protocols.

| sequence of indenter diameter | anatomical plane | gray to white matter stiffness ratio (-) |  |  |  |
| --- | --- | --- | --- | --- | --- |
|  |  | D20 | D100 | D200 | D500 |
| PA: 20 → 100 → 500 | transverse | 0.87 ± 0.27 | 1.09 ± 0.24 | - | 0.72 ± 0.13 |
| PB: 500 → 100 → 20 | transverse | 0.84 ± 0.38 | 1.10 ± 0.42 | - | 0.85 ± 0.48 |
| PC: 100 → 200 → 500 | transverse | - | 1.03 ± 0.36 | 1.06 ± 0.33 | 0.71 ± 0.24 |
| PD: 100 → 200 → 500 | coronal | - | 0.99 ± 0.21 | 0.99 ± 0.38 | 1.14 ± 0.59 |

Colors indicate ratios as follows: turquoise, ratios smaller than 1; white, ratios equal to 1; magenta, ratios greater than 1.

In Protocol B (Fig. 2b, f, j, n), we have reversed the order of applied indenter – from largest to smallest diameters vs. from smallest to largest for Protocol A. The results of Protocol B show a decrease in gray matter stiffness by a factor of 0.4 (*P* = 6.1 · 10^−5^) from the smallest to the intermediate indenter diameter and a decrease by a factor of 0.6 (*P* = 3.8 · 10^−11^) from the intermediate to the largest indenter diameter (Fig. 2b and Tab. S3, second row of upper table). The course of normalized gray matter stiffness across the different indenter diameters follows the same trend as for Protocol A, with an increase by 23% from the smallest to the intermediate and a decrease by 16% (*P* = 0.0047) from the intermediate to the largest indenter diameter (Fig. 2f and Tab. 2, second row). Similar to Protocol A, spinal cord white matter tissue stiffness (Tab. S3, second row of lower table) decreases by a factor of 0.3 (*P* = 1.3 · 10^−5^) from the smallest to the intermediate diameter and decrease by a factor of 0.9 (*P* = 0.020) from the intermediate to the largest diameter for Protocol B. This trend is also reflected in the results of normalized tissue stiffness (Fig. 2f and Tab. 2, second row) with a decrease by 15% from the smallest to the intermediate and an increase by 14% (*P* = 0.028) from the intermediate to the largest indenter diameter.

The sample-specific stiffness ratios (Fig. 2j) generally decreased from the intermediate to the largest diameter, except for Samples 13 and 16. In Samples 9, 10, 11, 15, and 16, gray matter consistently shows higher stiffness than white matter. Sample 12 demonstrated a continuous decrease in stiffness ratio with increasing diameter, whereas Sample 13 exhibited the inverse trend. In Samples 9, 10, 11, 14, and 15, stiffness ratios initially increased from the smallest to the intermediate diameter, followed by a decrease from the intermediate to the largest diameter. Samples 9, 10, 11, and 15 consistently showed stiffer gray matter compared to white matter, while Sample 13 showed the opposite trend. Additionally, stiffness ratios shifted from greater than one (stiffer gray matter) to smaller than one (stiffer white matter) between the smallest and intermediate diameters in Sample 16, and between the intermediate and largest diameters in Sample 12. The trend of the overall mean ratios averaged over all samples for each indenter diameter (Fig. 2n and Tab. 3, second row) is similar for Protocol B (reversed order) as for Protocol A (progressively increasing indenter diameter) with gray matter being stiffer than white matter for the intermediate diameter and softer for the smallest and largest diameter. A fictitious diameter of 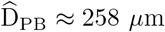 *µ*m can be estimated based on these data, at which the stiffness of spinal cord gray and white matter would be equal (ratio = 1). The slope of the connecting line is with a value of − 6 · 10^−4^ comparable to the corresponding slope of Protocol A.

To better resolve at which indenter size the shift in stiffness ratio happens, we refined the experimental design in Protocol C by excluding the 20 *µ*m diameter indenter and instead including a 200 *µ*m diameter indenter as the second-largest size. Since comparisons between Protocols A and B demonstrate that the order of indentations with different diameters does not affect the outcome, we adopted a progressively increasing indenter size sequence for Protocol C, following the approach of Protocol A, which had produced a higher number of valid indentations overall. The results of Protocol C show a decrease in gray matter stiffness by a factor of 0.7 (*P* = 1.6 · 10^−6^) from the intermediate to the second-largest indenter diameter and an increase by a factor of 1.4 (*P* = 6.7 · 10^−6^) from the second-largest to the largest indenter diameter (Fig. 2c and Tab. S3, third row of upper table). The course of the normalized gray matter stiffness across the different indenter diameters shows an increase by 1% from the intermediate to the second-largest and a decrease by 20% (*P* = 0.00098) from the second-largest to the largest indenter diameter (Fig. 2g and Tab. 2, third row). Similar to the results of Protocols A and B, spinal cord white matter tissue stiffness (Tab. S3, second row of lower table) decreases by a factor of 0.8 (*P* = 6.2 · 10^−6^) from the intermediate to the second-largest diameter and increases by a factor of 2.0 (*P* = 1.6 · 10^−17^) from the second-largest to the largest diameter for Protocol C. This trend is also reflected in the results of normalized tissue stiffness (Fig. 2g and Tab. 2, third row) with a an increase by 14% (*P* = 0.0098) from the second-largest to the largest indenter diameter.

Regarding the stiffness ratio between spinal cord gray and white matter, the results for Protocol C (Fig. 2k and o) show a consistent trend of decreasing stiffness ratio from the intermediate to the largest indenter diameter for Samples 18 and 20. When comparing only the results from indentations with the second-largest and largest diameters, the stiffness ratio decreases for all samples. We can observe a notable trend of increasing stiffness ratio from the intermediate to the second-largest indenter diameter explicitly displayed by Samples 17, 21 and 22. For Samples 19, 20, 21 and 24, gray matter is stiffer than white matter for the intermediate and the second-largest diameters. This dominance in stiffness shifts towards white matter tissue becoming stiffer than gray matter tissue in the majority of samples (except Samples 17 and 21) when indenting with the largest diameter. The course of overall mean ratios across all samples for each indenter diameter (Fig. 2o and Tab. 3, third row) underscores the observed trends for Protocols A and B with gray matter being stiffer than white matter for the intermediate and the second-largest diameters and softer for the largest diameter. A fictitious diameter of 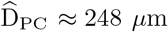 can be estimated based on these data, at which the stiffness of spinal cord gray and white matter would be equal (ratio = 1). Due to our refinement in the experimental setup, we achieve a better approximation for 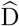 in Protocol C with a slope of the connecting line of − 11 · 10^−4^ which is substantially steeper than those observed in Protocols A and B.

Fig. 3 shows the shift in gray to white matter stiffness ratio for the combined datasets of Protocols A, B and C across the four different indenter diameters used in this study. This shift is observed between the 200 and the 500 *µ*m diameter with a slope of the connecting line of −10 · 10^−4^ and a fictitious diameter of 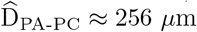, at which the stiffness of spinal cord gray and white matter would be equal (ratio = 1).

**Figure 3:**
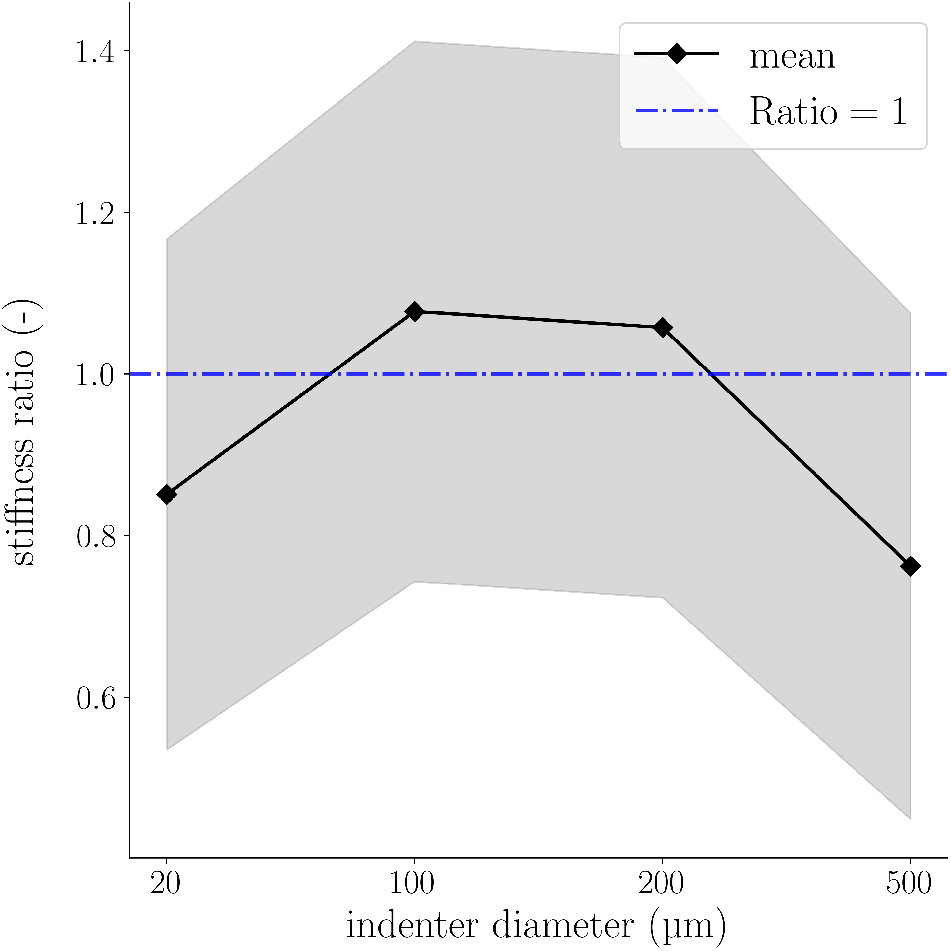
Ratio of spinal cord gray to white matter stiffness shifts between indenter diameters of 200 and 500 µm across Protocols. Mean ratio values of all biological replicates across different indenter diameters and Protocols A to C in black with standard deviation in gray. The blue dashed line indicates a ratio of one.

The observed reversal in the stiffness ratio between gray and white matter with increasing indenter size may reflect interactions between the probing scale and characteristic microstructural features of spinal cord tissue, such as axonal fiber organization and the heterogeneous distribution of cellular and extracellular components. This observation indicates that spherical indenters of different sizes mechanically activate spinal cord tissue in distinct ways and that gray and white matter, due to their inherent differences in microarchitecture, may exhibit opposing trends in their stiffness profiles across different indenter sizes.

### 2.4. Anatomical plane-dependent stiffness values and ratios of spinal cord gray to white matter tissue

To investigate whether the observed trends depend on the anatomical plane on which indentations are performed, we repeated Protocol C at the coronal plane in Protocol D. The results of Protocol D show a decrease in gray matter stiffness by a factor of 0.85 (*P* = 0.008) from the intermediate to the second-largest indenter diameter and an increase by a factor of 1.4 (*P <* 0.0001) from the second-largest to the largest indenter diameter (illustration in Fig. 2d and Tab. S3, last row of upper table). Contrary to the results for Protocols A to C, for the normalized apparent moduli, we observe no significant changes across different indenter diameters (Fig. 2h and Tab. 2, last row). Spinal cord white matter tissue stiffness (Tab. S3, last row of lower table) decreases by a factor of 0.86 (*P* = 0.0288) from the intermediate to the second-largest diameter and increases by a factor of 1.6 (*P <* 0.0001) from the second-largest to the largest diameter. Similar to gray matter tissue, we observe no change in normalized apparent moduli for the coronal plane of white matter tissue across different indenter diameters (Fig. 2h and Tab. 2, last row).

Regarding the stiffness ratio, Samples 25 to 32 (Fig. 2l) show no clear trend: Samples 26 and 32 show no significant change in stiffness ratio over the different diameters. The other samples show a mix of increasing and decreasing stiffness ratios from the intermediate to the second-largest indenter diameter and from the second-largest to the largest indenter diameter. Samples 28, 31 and 32 have consistently higher mean apparent moduli in gray matter than in white matter tissue. Samples 27, 29, and 30 show the opposite trend. Sample 25 sticks out of the dataset with significantly higher stiffness in spinal cord white matter tissue for indentations using the largest diameter. This also impacts the overall mean trend (Fig. 2p) to increase from the second-largest indenter diameter to the largest diameter. The mean ratios averaged over all samples for each indenter diameter in Protocol D performed on the coronal plane (Tab. 3, last row) show a different trend than Protocols A to C performed on the transverse plane: there is no clear shift in the ratio, which is approximately one (gray matter equals white matter stiffness) for the intermediate and the second-largest indenter diameter and slightly larger than one (gray matter stiffer than white matter) for the largest indenter diameter.

The different behavior observed between tests performed in the transverse and coronal planes may be related to the anisotropic architecture of spinal cord tissue, particularly the preferential orientation of axonal fibers relative to the indentation direction, especially in white matter. This suggests that sample orientation influences which structural components contribute most strongly to the measured mechanical response, an observation that is especially relevant in the context of the biphasic nature of CNS tissue.

## 3. Discussion

To the best of our knowledge, this is the first study to perform consecutive indentation tests on identical CNS tissue regions with varying indenter diameters, enabling a controlled assessment of sizeand planedependent variations in gray and white matter stiffness values and their ratio. While we confirmed typical inter-sample variability of CNS tissue under indentation [11] (statistical analysis: Tab. S8 to S47), we observed distinct trends related to changes in indenter diameter and anatomical plane. Our results show that CNS tissue stiffness measures are affected by indenter size. The mechanical response of gray matter tissue is uniformly increasing in normalized stiffness from the smallest (20 *µ*m diameter) to the intermediate indenter diameter (100 *µ*m diameter) and decreasing from the intermediate to the largest indenter diameter (500 *µ*m diameter) for all protocols with indentations on the transverse plane (Protocols A to C, Tab. S4 and Tab. 2). The response of white matter tissue shows the opposite trend: the stiffness decreases from the smallest to the intermediate indenter diameter (intermediate to the second-largest, 200 *µ*m diameter, for Protocol C), and then increases from the intermediate diameter (second-largest for Protocol C) to the largest diameter for tests on the transverse plane. Interestingly, for tests on the coronal plane, the normalized mechanical properties of CNS tissue are independent of the size of the indenter diameter for both gray and white matter tissue (Tab. 2, last row).

Compared with previously published indentation studies on spinal cord tissue, the present results follow the same general scale-dependent pattern, but yield overall higher stiffness values. In our data, gray matter stiffness ranged from approximately 530 to 3430 Pa and white matter stiffness from 560 to 4400 Pa, depending on protocol, anatomical plane, and indenter diameter. By comparison, studies using smaller AFM-based spherical probes generally reported markedly lower values, including 177 Pa (white) and 420 Pa (gray) for an 89.3 *µ*m indenter in rat spinal cord ([13]), 200 Pa (white) and 450 Pa (gray) for an 80 *µ*m indenter ([14]), 62.92 Pa (white) and 29.82 Pa (gray) for a 37.3 *µ*m indenter in zebrafish spinal cord ([15]), and 404 Pa (white) and 405 Pa (gray) for a 5 *µ*m indenter in mouse spinal cord ([20]). In contrast, the large-scale porcine spinal cord measurements of Bailly et al. ([11]), performed with a 500 *µ*m flat punch, yielded 510 Pa in white matter and 530 Pa in gray matter, which are closer to the lower end of the values reported here, particularly for our coronal measurements and some results on the transverse planes for an indenter with 500 *µ*m bead diameter. Taken together, these comparisons suggest that the present study extends the trend already visible in the literature: with increasing indenter size, the measured response shifts from a more local mechanical probing of individual microstructural components toward a more collective tissue-level response. At the same time, the fact that our values remain higher than most previous reports likely also reflects methodological differences beyond indenter diameter alone, including species, sample preparation, probe geometry, loading protocol, and the specific modulus definition used for analysis. Most importantly, however, the literature and the present results consistently show that both the absolute stiffness values and the white-to-gray matter relationship depend strongly on the spatial scale at which the tissue is mechanically probed.

While this is the first study to systematically investigate how varying indenter sizes affect the mechanical properties of identical CNS tissue locations, previous works have employed different (larger) indenter sizes on the entire spinal cord organ with the pia-arachnoid construct intact [31]. Therefore, Jin et al. indented directly on the pia–arachnoid construct. Thereby, they mechanically loaded the underlying spinal cord tissue through this outer membrane. They found decreasing stiffness with increasing indenter sizes. This observed behavior likely arises from the increasing mechanical activation of the significantly softer underlying spinal cord tissue for larger indenter sizes and the interplay between the pia-arachnoid complex and the underlying tissue in general. Extending the comparison to other biological tissues, Oyen et al. found that bone specimens exhibit nearly identical shear moduli when sequentially indented with a 1.63 mm diameter steel sphere and a 566 *µ*m diameter sapphire sphere [24]. The ability of bone to maintain a uniform response across indenter sizes underscores how mechanically complex – and scale-dependent – CNS tissue is. Wahlsten et al. reported an interesting approach, where they tested human skin tissue with AFM (6.10 *µ*m diameter), microindentation (100 *µ*m radius), and tissue-scale tensile loadings [26]. They report apparent moduli of 0.1 to 10 kPa for AFM and microindentation tests and elastic moduli of 100 to 200 kPa for unaxial tensile testing. To investigate the mechanisms behind their experimental observations, the authors built an *in silico* composite model, in which a collagen-fiber network is embedded in a ground matrix. The simulations reveal that the strength of the fiber–matrix bond dictates the scale dependence of the response: the fiber network contributes during indentation with the larger indenter only when this bond is strong enough to activate the fiber network and thereby enables effective load transfer between phases. These findings emphasize that capturing microstructural heterogeneity is critical for material models aiming to reproduce experimental behavior across spatial scales.

Since gray and white matter show different scale-dependent effects, their stiffness ratio may shift for changing indenter sizes: In general, gray matter is stiffer than white matter for small and softer for large indenters. For the first time, using the same setup and identical tissue locations, we observed a shift from gray matter being stiffer at smaller indenter sizes to white matter being stiffer at larger indenter sizes. Consequently, while prior research on spinal cord [31] or brain tissue [7, 32, 21] using differing indenter sizes had produced inconclusive findings regarding the ratio of CNS gray and white matter tissue stiffness values, the experimental protocols presented here successfully reconcile some of the previously reported discrepancies. Based on our observations, including the differences between stiffness values measured on transverse and coronal planes, we propose two possible explanations for the size-dependent phenomenon.

One factor contributing to the observed trends may be the anisotropy in diffusion in spinal cord tissue (see Tab. S7), as illustrated in Fig. 4a: for white matter, the diffusivity along axons (*λ*_∥_) is remarkably higher than across axons (*λ*_⊥_), whereas gray matter displays the same trend but to a smaller degree (*λ*_⊥,wm_ ≪ *λ*_∥,wm_; *λ*_⊥,gm_ < *λ*_∥,gm_). During transverse-plane indentations (Protocols A–C) fluid escapes mainly along the loading axis – which in white matter coincides with the axonal direction – while radial outflow is more obstructed. This impeded outflow pressurizes the interstitial fluid and results in an increase in measured stiffness. This mechanism is also present in gray matter, but attenuated by the weaker diffusion anisotropy and the larger extracellular space. Increasing indenter diameters trap a greater fluid volume, amplifying pressurization; because anisotropy is most pronounced in white matter, its stiffness rises more steeply with indenter diameter than that of gray matter, driving the observed shift in their stiffness ratio on the transverse plane. For coronal plane indentations (Protocol D), by contrast, radial outflow is largely unobstructed and now partly aligns with the axons. Pressurization therefore builds up only weakly as the indenter diameter increases, so that white matter stiffening is minimal and the more isotropic gray matter behavior [33] only shows minor changes. The result is an indenter-size-independent gray-to-white matter stiffness ratio on the coronal plane.

**Figure 4:**
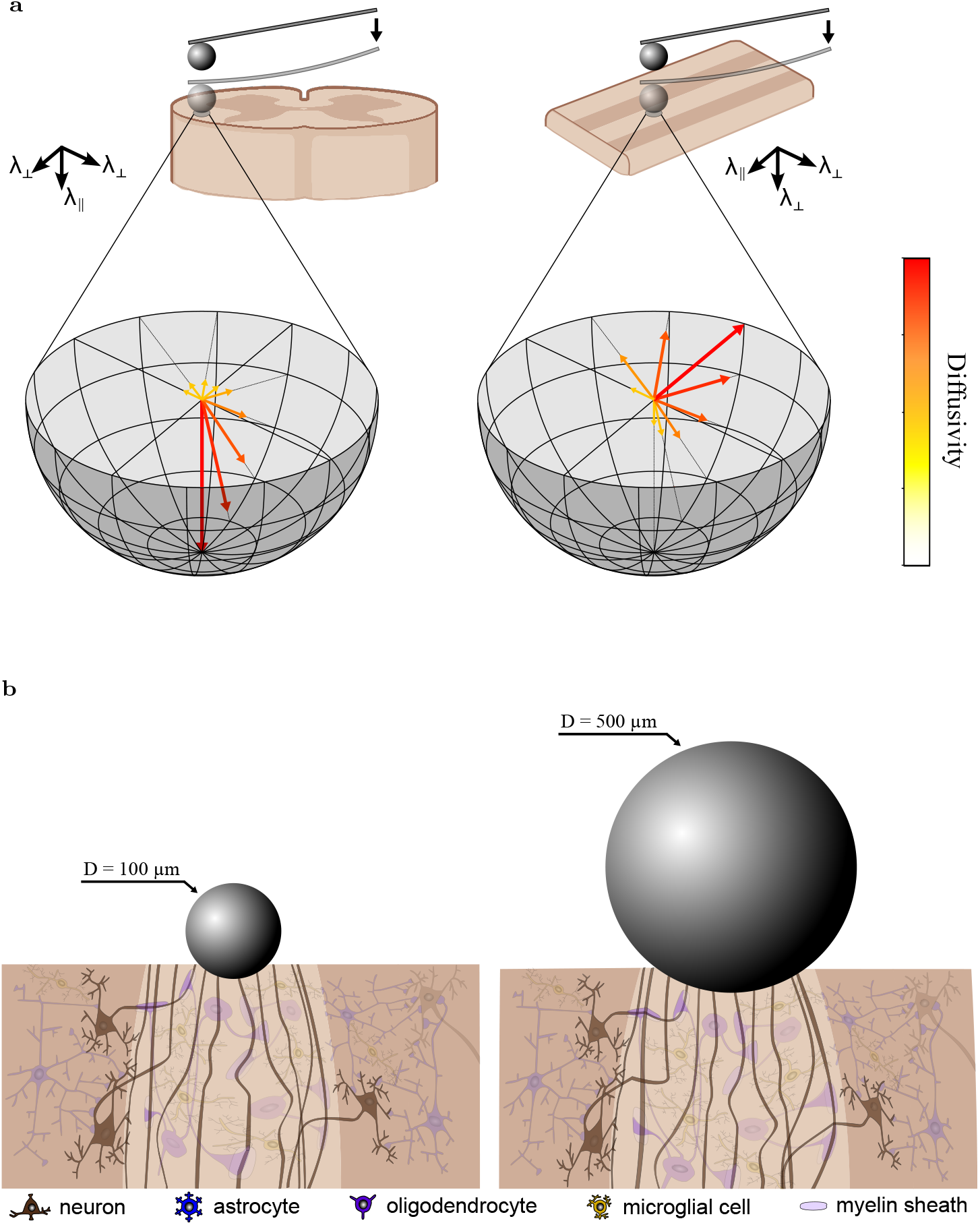
Diffusion anisotropy and CNS tissue microstructural architecture explain shifts in stiffness ratio of gray to white matter stiffness. (**a**) Schematic highlighting diffusion anisotropy of spinal cord white matter during spherical indentation on the transverse and coronal planes; arrows and colours depict diffusivity. (**b**) Schematic of deformation state during spherical indentation. Spinal cord white matter tissue is shown with axonal tracts, myelinating oligodendrocytes and microglia; myelin sheaths are depicted only exemplarily on selected axons. Spinal cord gray matter is shown with neurons, protoplasmic astrocytes and microglia cells. Both schematics are illustrative and not to scale. The spinal cord sketch was created using BioRender.

A complementary interpretation of our findings, particularly from Protocols C and D, highlights structural differences between gray and white matter associated with their distinct microarchitectures. White matter in the spinal cord mainly consists of myelinated, cranio-caudally orientated axons with an average diameter of 5 *µ*m, showing an anisotropic structure [33], whereas gray matter is comparatively isotropic [34, 33]. Although Feng et al. found white matter to be stiffer when indented perpendicular rather than parallel to the axons, they used a 19.1 by 1.6 mm rectangular flat-punch, which differs fundamentally from the spherical probes employed here [33]. Therefore, the absolute moduli cannot be compared directly. We assume that the measured response of spinal cord tissue is largely controlled by the underlying microstructural architecture. In white matter, the uniaxially oriented axons are closely connected to the network of glial cells through the myelin sheaths. Consequently, indenting spinal cord white matter with a 500 *µ*m diameter bead yields a relatively stiffer response than with a 100 *µ*m diameter, as the whole network is loaded instead of only individual components. In addition, the dimensions of the cellular components in our spinal cord samples in relation to the size of the indenter may also have affected the quality of these measurements. With cell body diameters of 5-10 and 6-8 *µ*m for astrocytes [35] and oligodendrocytes [36] in white matter tissue and up 40-60 *µ*m in diameter for neurons [37] in gray matter tissue, it remains debatable if tests with a 20 *µ*m diameter may have tested different types of cellular components in the spinal cord samples individually rather than a mechanical tissue response. Fig. 4b shows a schematic visualization of this assumed mechanism of scale dependent load transfer. Other studies could show that myelin and cellular coupling of axons via the glia matrix play a crucial role during tensile mechanical loads [38]. The influence of microstructural kinematics in white matter tissue is also confirmed during axonal damage threshold investigations [39]. We hypothesize that this bond could also account for different mechanical properties of spinal cord white and gray matter tissue during indentation tests, similar to the computational findings from Wahlsten et al. [26]. To test this hypothesis, we conducted additional experiments on the coronal plane of the spinal cord samples. This way, we were able to indent perpendicular to the main axon orientation. The results (Fig. 2l and p) clearly confirm this hypothesis. No significant changes between the stiffness of spinal cord white and gray matter tissue tested with differently sized beads can be observed for this test. Following this rationale, indentations performed perpendicular to the main axonal fiber direction, as in tests on the coronal sections of spinal cord white matter, would be expected to activate the tissue network differently than indentations on transverse sections. In the coronal plane, axons are arranged such that even smaller indenters likely interact with multiple adjacent fibers simultaneously, because these structures are effectively stacked horizontally relative to the perpendicular loading direction. As a result, the difference between small and large indenters in terms of the number of mechanically engaged axons may be markedly reduced compared with transverse sections. This would lead to a more similar contribution of the axon–glia network across indenter sizes and could explain why no size-dependent reversal between white and gray matter stiffness was observed in the coronal plane. In addition, these findings suggest that the mechanical response of the axon-glia network is anisotropic, with load transfer mediated by the network being more pronounced when loading occurs along the dominant fiber direction than when it occurs perpendicular to it.

Both hypotheses should be tested histologically to gather more data on the microstructure of the samples, and computationally to systematically test certain underlying assumptions *in silico*. A comprehensive strategy that includes an appropriate experimental testing protocol, histologically inspired physical modeling and an appropriate computational simulation method, such as the finite element method, will allow for a more thorough understanding of the intricate mechanical signature of spinal cord tissue across different spatial scales and the tissue mechanics of the central nervous system in general. [40].

CNS tissue is stiffer in transverse sections than in coronal sections for diameters between 100 and 500 *µ*m. When comparing results across anatomical planes for the same indenter diameter (Protocol C: transverse vs. Protocol D: coronal), spinal cord gray matter consistently exhibits stiffer mechanical properties in absolute terms, with 1.4 to 1.7-fold stiffer response on the transverse plane across all indenter diameters. Spinal cord white matter exhibits a comparable increase in stiffness, with a 1.5 to 1.9-fold stiffer response on the transverse plane. We attribute this discrepancy to the biphasic mechanical behavior of CNS tissue [1]. Poroviscoelastic simulations of flat-punch indentation indicate that interstitial fluid escapes along paths of least hydraulic resistance [41]. On the transverse surface, these paths are confined largely to the loading axis, leaving little space for fluid to drain, slowing its egress, and augmenting load support by pressurized fluid; the tissue therefore appears stiffer. In the coronal orientation, by contrast, fluid can migrate in circumferential direction – perpendicular to the indentation axis – into a much larger pore volume, facilitating rapid outflow, lowering fluid pressure and making the tissue appear more compliant.

Interestingly, the results reported by Koser et al. [17] show the opposite trend, where spinal cord white matter tissue was tested to be stiffer on coronal and sagittal sections compared to transverse sections, while for spinal cord gray matter tissue, no significant differences across different anatomical planes were observed. These discrepancies could be related to differences in the indenter diameter, as Koser et al. performed atomic force microscopy indentation tests using a 37.28 *µ*m diameter [17], while our smallest diameter for the coronal sections was 100 *µ*m in diameter: The smaller indenters may rather load individual components, while larger indenters load the entire network of cells and extracellular matrix. Again, the interplay between variations in the spatial regime of the mechanical loading (here the indenter diameter) and the architectural heterogeneity of the CNS tissue microstructure most likely modulate the observed mechanical properties on different anatomical planes and spatial scales.

It should be noted that some samples in this study are represented by comparatively low numbers of successful tests. This resulted from aspects of the experimental procedure, including the requirement to change the indenter tip twice for each sample, as well as limitations imposed by the surface topology of certain specimens – particularly in tests using the smallest indenter with a diameter of 20 *µ*m. Under these conditions, obtaining a uniformly high number of successful measurements across all samples was not always feasible. Consequently, several samples are associated with relatively small sample sizes. We acknowledge that the statistical robustness of these datasets is therefore lower compared to samples with larger numbers of successful tests, and the corresponding results should be interpreted with this limitation in mind. Importantly, this limitation applies only to a small number of samples – primarily those measured with the smallest indenter diameter – and we therefore believe that the observed trends and their interpretation remain valid.

We acknowledge that the clinical relevance of the presented results should be considered in light of potential differences between porcine and human spinal cord tissue. In this context, it is important to note that porcine spinal cords are widely used as translational models due to their morphological and physiological similarity to humans [42]. In particular, they closely resemble the human spinal cord in terms of size, dimensions, vertebral body height, circulatory system, lumbosacral enlargement length, cross-sectional area, and gray matter morphology [43, 44, 45].

Furthermore, the mechanical properties of porcine spinal cord and brain tissue have been shown to differ between *in vivo* and *ex vivo* measurements across various mechanical testing modalities [46, 47, 48]. In this study, all tests were conducted on *ex vivo* tissue to ensure comparability with the previously mentioned, partly contradictory results reported in the literature and due to the technical challenges associated with performing indentation tests *in vivo*. Therefore, this study does not address whether the observed trends are also present *in vivo*.

This study primarily focused on elucidating changes in measured mechanical properties based on indentation data analyzed using the conventionally employed linear Hertz model. Future studies could employ nonlinear continuum mechanical models to incorporate nonlinear effects in the analysis. As mentioned in the Methods section, a Poisson’s ratio of 0.5 was used in this study. We do not claim that spinal cord tissue behaves strictly incompressible. Rather, this assumption was adopted to ensure comparability with the majority of existing indentation studies and to maintain a commonly used modeling framework. Nevertheless, we expect that nonlinear continuum mechanical models employing slightly compressible material parameters would yield similar trends to those reported here. Validation of this assumption should be addressed in future studies.

Because measurements with different indenter diameters were performed at the same locations within each sample, repeated indentation could in principle introduce history-dependent effects, such as localized microstructural damage or fluid redistribution. However, preliminary tests involving repeated indentations at the same location with substantially shorter time intervals did not reveal systematic changes in the measured response. Furthermore, the time between consecutive measurements in the present study typically exceeded one minute due to the required indenter recalibration steps. Therefore, we consider it unlikely that repeated loading substantially influenced the observed trends.

All measurements in the present study were conducted using a single indentation rate of 50 *µ*m/s and the reported material parameters were derived from the loading portion of the force–indentation curves without including a holding phase for stress relaxation. As nervous tissue exhibits pronounced viscoelastic and potentially poro-elastic behavior, time-dependent processes, such as fluid redistribution and pressure dissipation, may influence the measured mechanical response. Consequently, the parameters reported here should be interpreted as effective elastic moduli characterizing the loading response rather than equilibrium properties. Future studies incorporating controlled hold phases and relaxation analysis across different time scales would allow a more detailed investigation of fluid pressurization effects and their potential contribution to the scale-dependent mechanical behavior observed in this work.

Previous indentation-based studies [4, 12] further suggest that increasing temperature affects the absolute apparent stiffness of both gray and white matter similarly, with little influence on their stiffness ratio.

The investigated indenter size range was limited to spherical beads with diameters of 20, 100, 200, and 500 *µ*m. While these sizes cover several relevant spatial scales for indentation-based characterization of spinal cord tissue, additional indenter sizes – especially above 1 mm diameter – could further refine the understanding of how measured mechanical properties depend on the spatial scale of probing. Additionally, the applied strain range during indentation was limited by the selected testing protocol. Future studies should investigate whether the reported trends persist across a broader range of strains. Lastly, all measurements were conducted at a single indentation rate of 50 *µ*m/s. Since nervous tissue exhibits viscoelastic behavior, different loading rates may influence the measured mechanical response. However, the comprehensive study by Becker et al. demonstrated that variations in measurement parameters, including loading rate, affect the absolute elastic moduli of gray and white matter but cannot fully reproduce the shifts in the gray-to-white matter stiffness ratio [4] observed across different indenter sizes. Therefore, we assume that the trends observed in the present study are primarily scale-dependent rather than a consequence of the viscoelastic behavior of spinal cord tissue. Still, we acknowledge the findings of Becker et al. [4] and cannot quantitatively exclude that the magnitude of the trends observed in the present study may vary for different choices of loading rate or maximum indentation depth. Further experimental studies are required to investigate a broader testing parameter space.

In conclusion, we show that the long-standing question of whether white matter is stiffer or more compliant than gray matter can be resolved by accounting for the spatial scale of indentation. Systematically varying spherical indenter size, we identify the diameter range over which the gray to white matter stiffness ratio reverses – from > 1 (gray stiffer) to < 1 (white stiffer) – and confirm that this reversal is independent of indentation order. This shift in stiffness is observed for indenter diameters between 200 and 300 *µ*m. On the transverse plane (parallel to axons), increasing indenter size yields a markedly stiffer white matter response and a modest gray matter stiffening, whereas on the coronal plane (perpendicular to axons) the stiffness ratio remains invariant across diameters. These results provide critical benchmarks for continuum mechanical models of spinal cord tissue interested in incorporating microstructural features (e.g., porosity, axonal alignment) and offer a dataset for their validation, advancing our mechanistic understanding of CNS tissue mechanics.

## Methods

### Sample handling

The sample preparation in this study is based on the same protocol described in [49]. We collected porcine spinal cord tissue from three to four months old female pigs from a local abattoir (Contifleisch GmbH und Unifleisch GmbH, Erlangen, DE) and embedded approximately 7 cm long segments in low melting point agarose (Carl Roth GmbH + Co. KG, Karlsruhe, DE). We prepared samples approximately 7 mm thick from these segments for subsequent indentation testing. During the preparation, we hydrated the samples with Dulbeccos phosphate buffered saline solution and kept the cutting plate cooled with ice packs to minimize tissue degradation. After preparation, we stored the agarose blocks refrigerated at 4 °C until they were tested. After testing, we applied two to three drops of Richardson’s Solution [50], consisting of 0.5% w/v methylene blue, 0.5% w/v azur II and 0.5% w/v di-sodium tetraborate decahydrate in distilled water. This amount of solution proved to be adequate to colour the samples to the desired intensity. Fig. 5 (**c**) displays a sample stained after application of the solution. Applying this solution enabled us to confirm whether individual grid scans were actually located in gray or white matter tissue. This additional validation was necessary, because the distinction between spinal cord gray and white matter is not always possible without additional staining, particularly at the interface, as illustrated in Fig. 5 (**a**), (**b**), (**d**), and (**e**). The *post mortem* times of all 32 samples are listed in Tab. S2 with a mean *post mortem* time per sample of 5.7 ± 1.7 h and a maximum time after harvest of 8 hours and 54 minutes. All tests were conducted at 22 °C room temperature.

**Figure 5:**
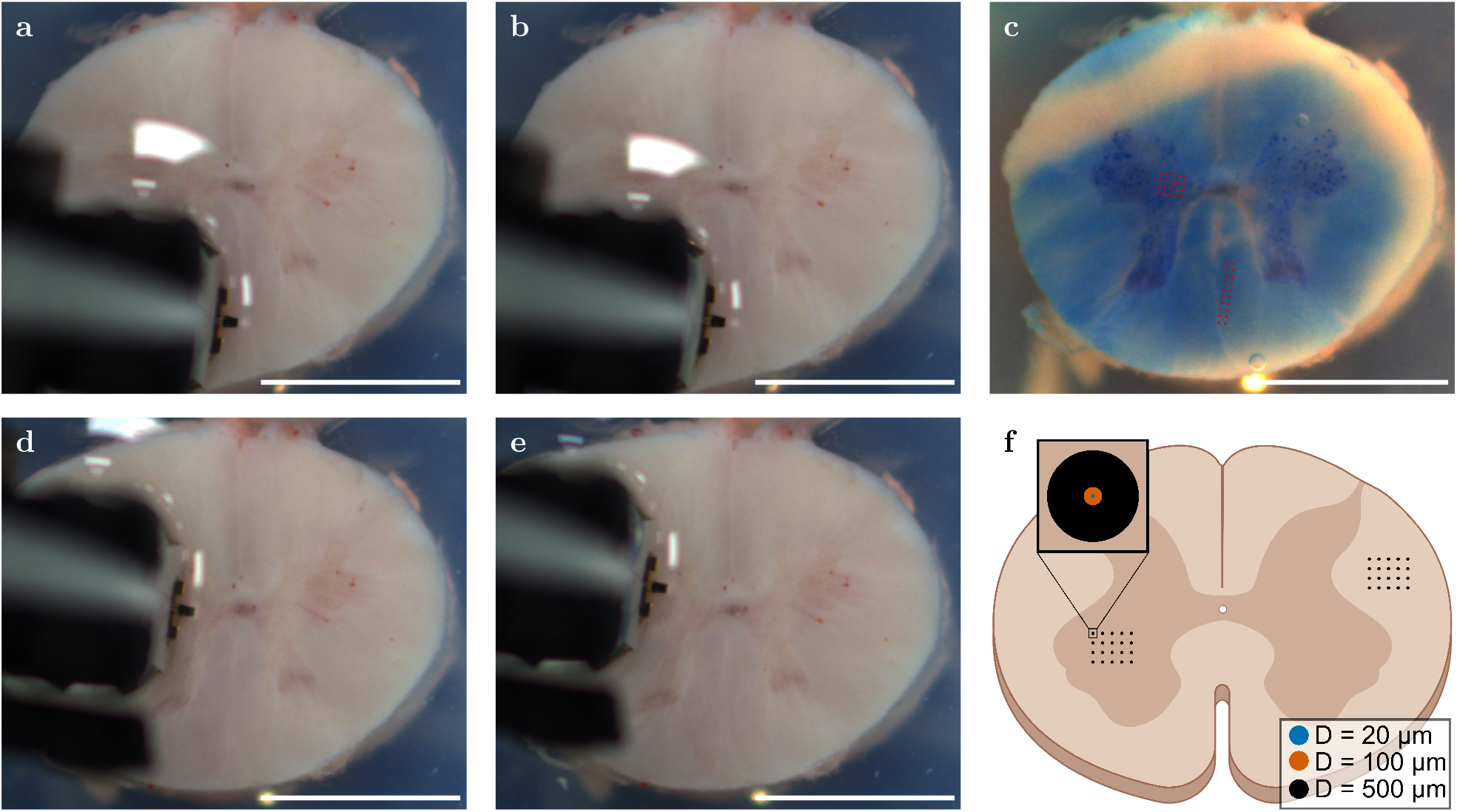
Allocation of indentation points on the transverse plane of a spinal cord slice. (**a**) and (**b**) show images of the start and end point of the 20-point grid scan on spinal cord white matter tissue, respectively. (**d**) and (**e**) show images of the start and end point of the 20-point grid scan on spinal cord gray matter tissue, respectively. (**c**) shows the same sample stained with Richardon’s solution. The two grid scans on white (two by ten) and gray matter (five by four) are indicated with red dots. (**f**) depicts a sketch to visualize the experimental methodology. Scale bars: 5 mm. The spinal cord sketches were created using BioRender.

### Experimental setup

We used a Chiaro Nanoindenter™ (Optics11 Life, Amsterdam, NL) for indentation experiments, which uses a piezo-electric translator to move the tip vertically during an indentation. The force is calculated by measuring the deflection of the cantilever to which the indenter tip is glued. As described in [49], the deflection is assessed by the phase shift between light that is reflected at the optical fiber-to-gap interface and the gap-to-cantilever interface [51].

To image the transverse plane of the spinal cord samples during the experiments, we mounted an industrial camera (IDS Imaging Development Systems GmbH, Obersulm, DE) together with a lens with 25 mm focal length (RICOH, Neu-Isenburg, DE) at a fixed position 10 cm above the sample (exemplary photographs can be seen in Fig. 5 (**a**) to (**e**)).

### Indentation procedure

We performed all indentations in a indentation-controlled mode and employed trapezoidal loadingholding-unloading profiles. During indentation-controlled experiments, a predefined threshold is used to detect the sample surface in real time, based on the deflection of the cantilever. Once the threshold is reached, it serves as the reference point for applying the desired indentation depth. From the findings in our previous study [49], we assumed the ratio of spinal cord gray to white matter stiffness to be constant for temperatures ranging between room and body temperature. This finding was independently confirmed from Becker et al. [4] using Atomic Force Microscopy (AFM) on rodent spinal cord. Since we intended to investigate a possible indenter size dependence of this ratio, we performed all experiments at room temperature to reduce heat-induced tissue degradation.

We used spherical indenter tips with a nominal stiffness of 0.5 N/m and nominal diameters of 20, 100, 200 and 500 *µ*m to test the mechanical properties of gray and white matter on different spatial scales. All indentations were performed with a loading rate of 50 *µ*m/s. First, we performed two 20-point indentation grids, one in spinal cord gray and one in spinal cord white matter tissue with a nominal indenter diameter of 20 *µ*m. Subsequently, we tested the same grids with the same indentation locations using indenters with a nominal diameter of 100 and 500 *µ*m. In the following this experimental procedure is termed as Protocol A (PA). With Protocol A we tested eight samples from seven different spinal cords. We then repeated the test procedure in Protocol B (PB) for the reverse order on eight samples from eight additional spinal cords: We started with the 500 *µ*m diameter tip, followed by the 100 *µ*m diameter tip, and ended with the 20 *µ*m diameter tip. This allowed us to check whether reversing the sequence of indenter diameters affects the overall trends. Next, in Protocol C (PC), we followed the same procedure as in Protocol A with progressively increasing indenter diameters of 100, 200 and 500 *µ*m on eight samples from eight additional spinal cords. In contrast to Protocol A, the 20 *µ*m indenter was omitted and replaced by a 200 *µ*m indenter to more precisely examine the indenter-size range in which the transition in the gray-to-white matter stiffness ratio was expected to occur based on the results of Protocols A and B, while also incorporating an intermediate indenter size that was expected to yield a higher number of successful indentations per grid scan than the smallest indenter used previously. All tests in Protocol A, B and C were conducted at the transverse plane of the spinal cord samples. To test whether the anisotropic characteristics of CNS tissue influence the observed overall trends, in Protocol D (PD), we performed indentation experiments on eight samples from eight additional spinal cords with increasing indenter diameters of 100, 200 and 500 *µ*m at the coronal plane of the spinal cord samples. The indentation depths were 10, 10, 15, and 25 *µ*m for tests with indenter diameters of 20, 100, 200, and 500 *µ*m, respectively. Tab. 4 gives a summary of the different protocols employed in this study.

**Table 4:** Overview of experimental indentation protocols.

| experimental protocol | anatomical plane | sequence of indenter diameter (in $\mu\text{m}$) |
| --- | --- | --- |
| Protocol A (Fig. S3a) | transverse | 20 → 100 → 500 |
| Protocol B (Fig. S3b) | transverse | 500 → 100 → 20 |
| Protocol C (Fig. S3c) | transverse | 100 → 200 → 500 |
| Protocol D (Fig. S3d) | coronal | 100 → 200 → 500 |

The mean experimental duration per sample was approximately 53 ± 14 min with a maximum duration of 91 minutes. In addition, we performed control experiments on 3 % agarose to validate the testing method, and, as expected for this homogeneous material, observed no indenter-size-dependent differences in stiffness (see Fig. S4)

### Indentation data analysis

We employ the same contact-point identification method as described in [49] and manually exclude bad curves that do not show the typical force-displacement curves expected for indentation experiments. For reproducibility, the raw data, including an overview of which curves were excluded during filtering, are publicly available on Zenodo (see Data availability). Fig. 5 illustrates our indentation grid allocation routine. We manually extract the start and end points from the images which were recorded during the experiment (see Fig. 5a to d) and superimpose these markers on the reference image of the stained sample (Fig. 5c). All the images, the start and end point of grid scans as well as the stained sample, were taken from the very same angle and distance to the sample. Using the predefined step sizes for the individual grid scans in x and y direction, we can now visualize the grid on the image of the stained sample (red indications in Fig. 5c) to ensure that the measurements were performed on either spinal cord gray or white matter tissue.

### Statistical analysis

A biological replicate is one spinal cord slice from an independent animal *n* = 32 animals. A technical replicate is a single indentation measurement. Each slice received 20 gray matter and 20 white matter indentations per indenter diameter (three different diameters per protocol, see Tab. 4).

A linear mixed-effects model was fit with tissue type (gray, white matter) and indenter diameter (20 *µ*m, 100 *µ*m, 500 *µ*m for Protocol A and B; 100 *µ*m, 200 *µ*m, 500 *µ*m for Protocol C and D) as fixed effects, and biological replicate as a random effect. Pairwise differences between diameters within each tissue were assessed on the technical replicates by unpaired two-sided Welch’s t-tests with Holm correction; raw and adjusted p-values, mean differences, and all test statistics are reported in the Supplementary Material. Additionally, we compared gray versus white matter at each diameter using Welch’s t-tests on the technical replicates (Holm-adjusted), and report those alongside Tukey’s honestly significant difference (HSD) test outcomes – mean differences, adjusted p-values, confidence intervals, and significance decisions (see Tab. S8 to S47).

### Mechanical properties

Following the approach presented in [49], we employ contact mechanics based on Hertz’s theory [52] to obtain an apparent modulus from force-displacement data of the loading curve. Due to higher flexibility in terms of the validity of models with regards to the maximum indentation depth, we use the Sneddon model [53] based on Hertz’s theory and the works of Segedin [54]. The model is described by the following relation,

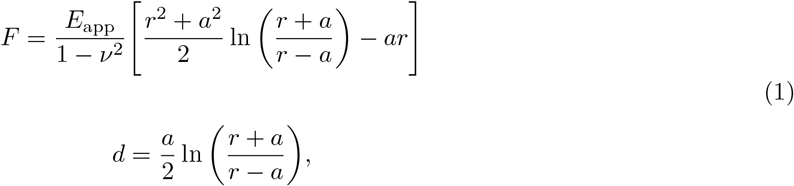

where *r* denotes the radius of the indenter, *F* is the measured force, *d* the displacement and *a* the radius of the circular contact area between indenter and sample to relate force to displacement. We assume a Poisson’s ratio of *ν* = 0.5 for the model fitting. The Poisson’s ratio was chosen to maximize comparability with literature data, as indentation-based fitting procedures traditionally assume an incompressible material behavior. In order to establish a consistent relation between indenter size and maximum indentation depth across different indenter diameters, we truncate the loading curves at 10% of the radius of the indenter prior to the fitting procedure. The Hertz model assumes a purely linear-elastic material response, which does not reflect the viscous and porous nature typically observed in brain and spinal cord tissue [55, 41, 56]. Consequently, in the following sections, *E*_app_ is treated as an approximate measure valid under particular loading conditions, primarily to compare ratios and identify trends. This apparent modulus should not be confused with more comprehensive mechanical parameters obtained from nonlinear continuum mechanics and inverse identification procedures [57], where one can computationally analyze the mechanical response of the spinal cord tissue with much greater flexibility, particularly with respect to the loading scale and the geometry of the indenter tip.

We evaluate the ratios of spinal cord gray matter to white matter stiffness in the following way: First, we compute the mean apparent modulus for each individual tissue type per indenter diameter and sample. With those values, we calculate the ratios per diameter for each sample by dividing the mean apparent modulus of all indentations on spinal cord gray matter tissue by the mean apparent modulus of all indentations on spinal cord white matter tissue. To illustrate the change in these ratios across the different indenter diameters we compute the mean and standard deviation of the ratios over all samples for each indenter diameter. Additionally, we use a normalization routine that reduces the influence of inter-sample variability and highlights only the trends in stiffness ratio changes across different indenter diameters. To normalize the results of each sample *S*_*p*_ (where *p* = 1, 2, …, *N*, and *N* is the total number of samples) at a given indenter diameter *D*_*q*_ ∈ {20, 100, 200, 500} *µ*m, we collect all apparent moduli for that sample-diameter combination into a set ℰ (*S*_*p*_, *D*_*q*_) and normalize each value by the mean of this set. Importantly, this normalization is performed using a common denominator for both spinal cord white matter and gray matter measurements within the respective (*S*_*p*_, *D*_*q*_) combination. Thus, each individual apparent modulus is expressed relative to the overall mean apparent modulus of all measurements obtained for that sample and indenter diameter, irrespective of tissue type. This approach provides a common sample-specific reference that preserves the relative differences between the two tissue types, while reducing the influence of inter-sample variability. In contrast, normalizing white matter and gray matter measurements separately by their respective tissuespecific means, would force both distributions to be centered around 1 and would therefore obscure how measurements obtained with different indenter diameters change within each tissue type relative to the overall response of the corresponding sample. Specifically:

- Let *n*_SD_ be the total number of measurements for the combination (*S*_*p*_, *D*_*q*_).
- We group the apparent moduli of these measurements into one set, ℰ (*S*_*p*_, *D*_*q*_).
- This set can be viewed as the union of the spinal cord white matter tissue (wm) and gray matter tissue (gm) subsets of measurements.

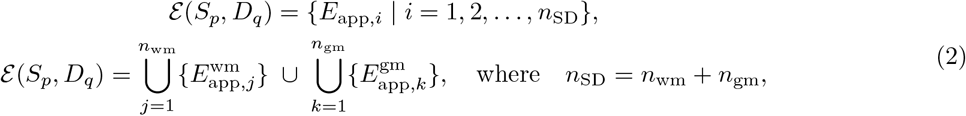

where *n*_wm_ and *n*_gm_ are the numbers of measurements taken in spinal cord white matter and gray matter tissue, respectively.
- Let ℰ^norm^(*S*_*p*_, *D*_*q*_) be the set of normalized apparent moduli, 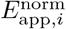, for the combination (*S*_*p*_, *D*_*q*_).

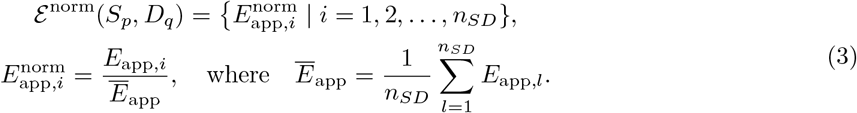

To compute overall mean values across all samples at a given diameter we define the subsets ℰ^wm^(*D*_*q*_) and ℰ^gm^(*D*_*q*_) for measurements on spinal cord white matter and gray matter tissue containing all apparent moduli of all samples *N* and one tip diameter *D*_*q*_. The mean values across samples for one tip diameter, 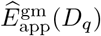 and 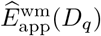, are derived as follows,

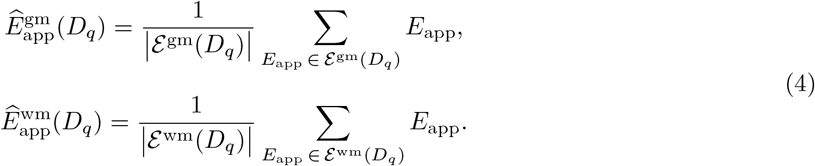

## Declaration of AI and AI-assisted technologies in the writing process

During the preparation of this work the author(s) used Deepl and ChatGPT in order to improve readability and language. After using this tool/service, the author(s) reviewed and edited the content as needed and take(s) full responsibility for the content of the publication.

## Data availability

The datasets generated and analyzed during the current study are publicly available in the Zenodo repository at: https://doi.org/10.5281/zenodo.19051452.

## Funding

This work was funded by the German Research Foundation (DFG) project 460333672 CRC1540 EBM (projects B01 and B04).

## Acknowledgements

We gratefully acknowledge the funding by the German Research Foundation (DFG) project 460333672 CRC1540 EBM (project B01 and B04).

## Author contributions

Conceptualization: O.N., S.B.; Formal analysis: O.N., S.B.; Investigation: O.N., H.V.S.; Methodology: O.N., T.F., M.H., S.B.; Project administration: S.B.; Resources: H.S.V, S.B.; Software: O.N.; Supervision: P.S., S.B.; Visualization: O.N. S.B.; Writing – original draft: O.N., S.B.; Writing – review and editing: O.N., M.H., S.K., T.F., R.G.R., P.S., S.B.

## Competing interests

The authors declare no competing interests.

## Notes

https://doi.org/10.5281/zenodo.19051452

### Competing Interest Statement

The authors have declared no competing interest.

