## Supplementary Material for "Spatial scale of indentation explains shift in ratio between spinal cord gray and white matter stiffness"

*Linear regression analysis*

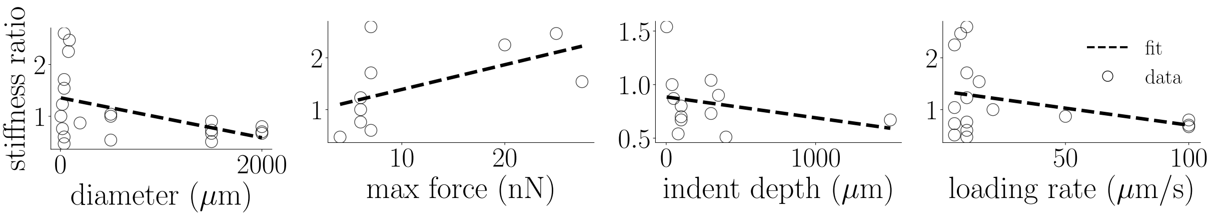


Fig. S1: **Overview of linear regression analyses** for possible correlations between the stiffness ratio of gray to white matter tissue and four different experimental parameters from the literature overview in Tab. S1. Correlation tests from left to right: stiffness ratio vs. indentation diameter, stiffness ratio vs. maximum force, stiffness ratio vs. maximum indentation depth, stiffness ratio vs. loading rate. The data points correspond to the values given in Tab. S1.

Table S1: **Literature review of indentation experiments on CNS tissue.**

| Year | Author | Species | Organ | Anatomical plane | Parameter | Indenter diameter ($\mu$m) | Indenter geometry | Temperature | Max. force or indentation depth | Loading rate | Model | White matter stiffness (Pa) | gray matter stiffness (Pa) | Ratio |
|---|---|---|---|---|---|---|---|---|---|---|---|---|---|---|
| 2010 | Van Dommelen et al. ([6]) | Pig | Cerebrum (Posterior) | Sagittal | Shear modulus | 2000 | Spherical | Room temperature | 100 $\mu$m | 100 $\mu$m/s | Lee and Radok ([58]) | 949 | 669 | 0.70 |
| | | | Cerebrum (Superior) | | | | | | | | | 1209 | 816 | 0.67 |
| | | | Cerebrum (Anterior) | | | | | | | | | 925 | 738 | 0.80 |
| 2015 | Budday et al. ([7]) | Cattle | Cerebrum | Transverse | Effective modulus | 1500 | Flat cylindrical punch | Room temperature | 300 $\mu$m | 5 $\mu$m/s | Mean slope method | 1895 | 1389 | 0.73 |
| 2016 | Weickenmeier et al. ([8]) | Cattle | Cerebrum | Sagittal | Effective modulus | 1500 | Flat cylindrical punch | Room temperature | 400 $\mu$m | 5 $\mu$m/s | Mean slope method | 1330 | 680 | 0.51 |
| 2018 | Weickenmeier et al.[a] ([9]) | Pig | Cerebrum | Transverse | Elastic shear modulus | 1500 | Flat cylindrical punch | Room temperature | 350 $\mu$m | N/A | Mean slope method | 300 | 270 | 0.90 |
| 2011 | Kaster et al.[b] ([10]) | Pig | Brain, exact location not reported | Not reported | Elastic modulus | 1500 | Flat cylindrical punch | Room temperature | 1500 $\mu$m, estimated strain 15–20% | 0.1 $s^{-1}$ | Samani method ([59]) | 1787 | 1195 | 0.67 |
| 2012 | Finan et al. [19] | Rat | Cerebellum | Sagittal | Long-term shear modulus | 500 | Flat cylindrical punch | "Warm" | 39.3 $\mu$m, estimated strain $\sim 10\%$ | N/A | Prony series expansion | 110 | 110 | 1.0 |
| 2022 | Sundaresh et al. ([5]) | Human | Cortex | Not reported | Maximum shear modulus | 500 | Flat cylindrical punch | Not reported | 40, 80, 120 $\mu$m, estimated strain $\sim 10, 20, 30\%$ | 1.9 $s^{-1}$ | Harding and Sneddon ([60]) | 1624 | 883 | 0.54 |
| 2025 | Bailly et al. ([11]) | Pig | Spinal Cord | Transverse | Effective Modulus | 500 | Flat cylindrical punch | 5 | 300 $\mu$m | 5 $\mu$m/s | Mean slope method | 510 | 530 | 1.04 |
| 2025 | Neumann et al. ([12]) | Pig | Spinal cord | Transverse | Apparent modulus | 195 | Spherical | Mean over 20 - 37 °C | 50 $\mu$m | 50 $\mu$m/s | Sneddon ([61]) | not applicable | not applicable | 0.87 |
| 2017 | Moeendarbary et al. ([13]) | Rat | Spinal cord | Transverse | Apparent modulus | 89.3 | Spherical (AFM) | Room temperature | 20, 30 nN | 5-10 $\mu$m/s | Hertz ([62]) | 177 | 420 | 2.47 |
| 2020 | Baumann et al. ([14]) | Rat | Spinal cord | Horizontal or Transverse (not specified) | Elastic modulus | 80 | Spherical (AFM) | 37 (°C) | 20 nN | 5 (°C) $\mu$m/s | Hertz ([62]) | 200 | 450 | 2.25 |
| 2020 | Möllmert et al. ([15]) | Zebrafish | Spinal cord | Transverse | Elastic modulus | 37.3 | Spherical (AFM) | 18 (°C) | 4 nN | 10 $\mu$m/s | Sneddon ([61]) | 62.92[c] | 29.82[c] | 0.47 |
| 2010 | Christ et al. ([16]) | Rat | Cerebellum | Sagittal | Apparent modulus | 37.28 | Spherical (AFM) | Room temperature | 3 $\mu$m, <27.5 nN | 15 $\mu$m/s | Hertz ([62]) | 294 | 454 | 1.54 |
| 2015 | Koser et al. ([17]) | Mouse | Spinal cord | Transverse | Apparent modulus | 37.28 | Spherical (AFM) | Not reported | 7 nN | 10 $\mu$m/s | Hertz ([62]) | 48 | 125 | 2.60 |
| | | | | Horizontal | | | | | | | | 75 | 128 | 1.71 |
| | | | | Sagittal | | | | | | | | 77 | 127 | 0.60 |
| 2020 | Eberle et al. ([18]) | Mouse (young) | Cerebellum | Sagittal | Apparent modulus | 20 | Spherical (AFM) | 18–23 °C | 6 nN | 10 $\mu$m/s | Sneddon ([61]) | 196.7 | 260.6 | 0.76 |
| | | Mouse (old) | | | | | | | | | | 221.9 | 273.1 | 1.23 |
| 2020 | Cooper et al. ([20]) | Mouse | Spinal cord | Transverse | Elastic modulus | 5 | Spherical (AFM) | Room temperature | 5-7 nN | 20.6 $\mu$m/s | Hertz ([62]) | 404 | 405 | 1.0 |

Summary sorted by indenter size in descending order. AFM, atomic force microscopy; [a]Single animal; [b]Finite-element derived; [c]Acquired from Fig. 3f of corresponding study using [63].

*Inter-sample variability for Protocol C and D*

**a**

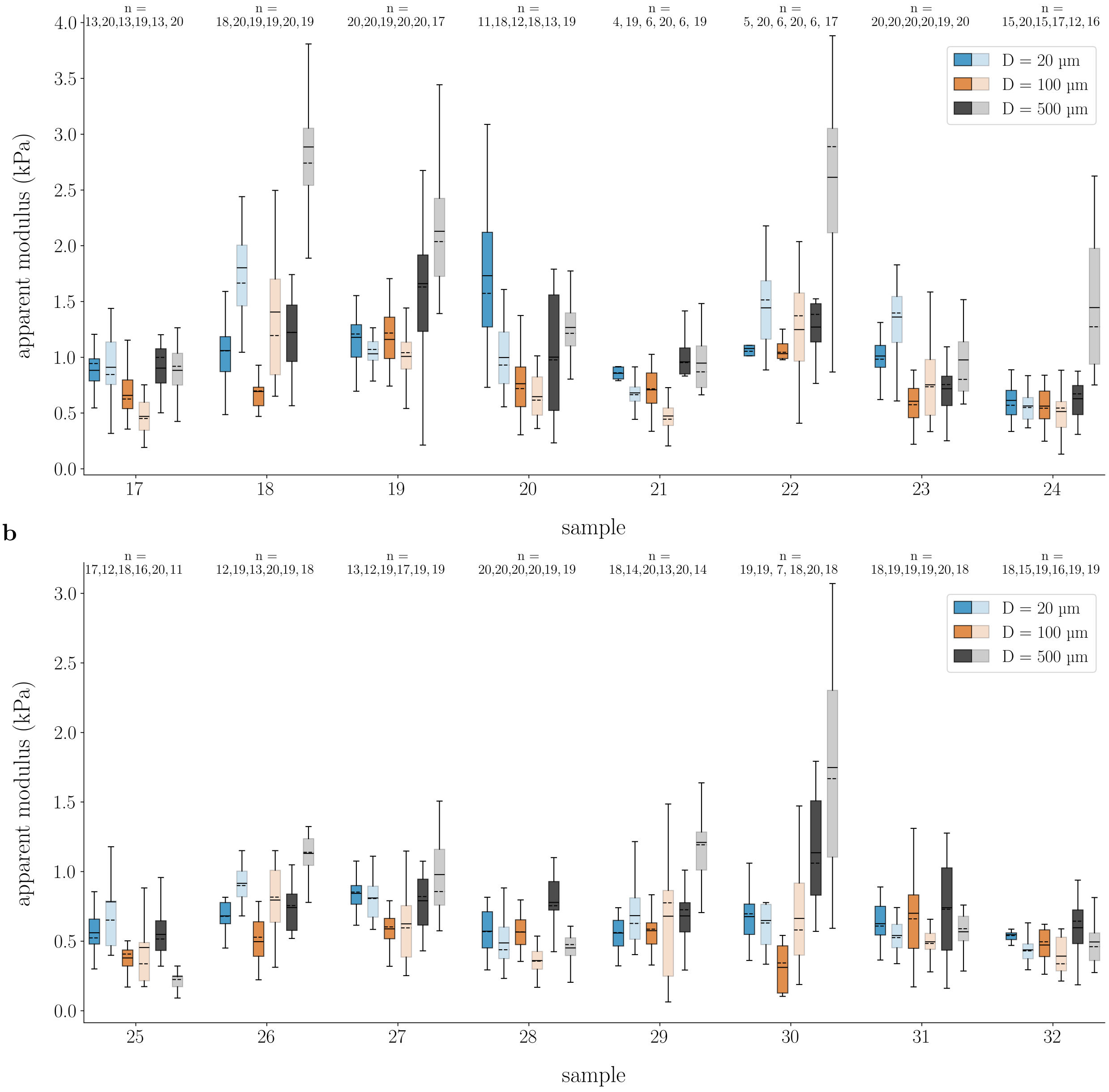


Fig. S2: **Apparent moduli for Protocol C and D indentations across samples for three different indenter sizes.** (**a**) Results for tests according to Protocol C, order of measurements: D = 100, D = 200, D = 500 $\mu$m), transverse plane. (**b**) Results for tests according Protocol D, order of measurements: D = 100, D = 200, D = 500 $\mu$m, coronal plane. Results are collectively clustered for each sample. Indentations with D = 100, 200 and 500 $\mu$m are shown in orange, bluish green and gray, respectively. Lighter colors depict results for spinal cord white matter tissue, darker colors for spinal cord gray matter tissue. Boxes show the 25th (Q1), 50th (median) and 75th (Q3) percentiles with inter-quartile range (IQR: from Q1 to Q3); whiskers reach the most extreme data points within $\pm$ 1.5 $\times$ IQR of Q1 and Q3. Outliers outside this range are not displayed but were included in the analysis. Data points are indicated as circles, mean values as continuous lines, and median values as dashed lines. n indicates the number of technical replicates.

*Sample preparation*

Table S2: ***Post mortem* times for biological replicates.**

| **sample** | ***post mortem* time (hh:mm)** |
|---|---|
| 1 | 4:11 |
| 2 | 7:29 |
| 3 | 5:42 |
| 4 | 8:54 |
| 5 | 4:39 |
| 6 | 8:00 |
| 7 | 3:43 |
| 8 | 7:49 |
| 9 | 3:11 |
| 10 | 5:33 |
| 11 | 7:21 |
| 12 | 7:04 |
| 13 | 4:38 |
| 14 | 6:20 |
| 15 | 7:50 |
| 16 | 4:00 |
| 17 | 7:12 |
| 18 | 3:33 |
| 19 | 3:55 |
| 20 | 6:00 |
| 21 | 7:46 |
| 22 | 3:33 |
| 23 | 5:39 |
| 24 | 6:49 |
| 25 | 4:05 |
| 26 | 6:02 |
| 27 | 7:24 |
| 28 | 3:50 |
| 29 | 5:34 |
| 30 | 4:37 |
| 31 | 3:35 |
| 32 | 6:57 |

*Indentation protocols*

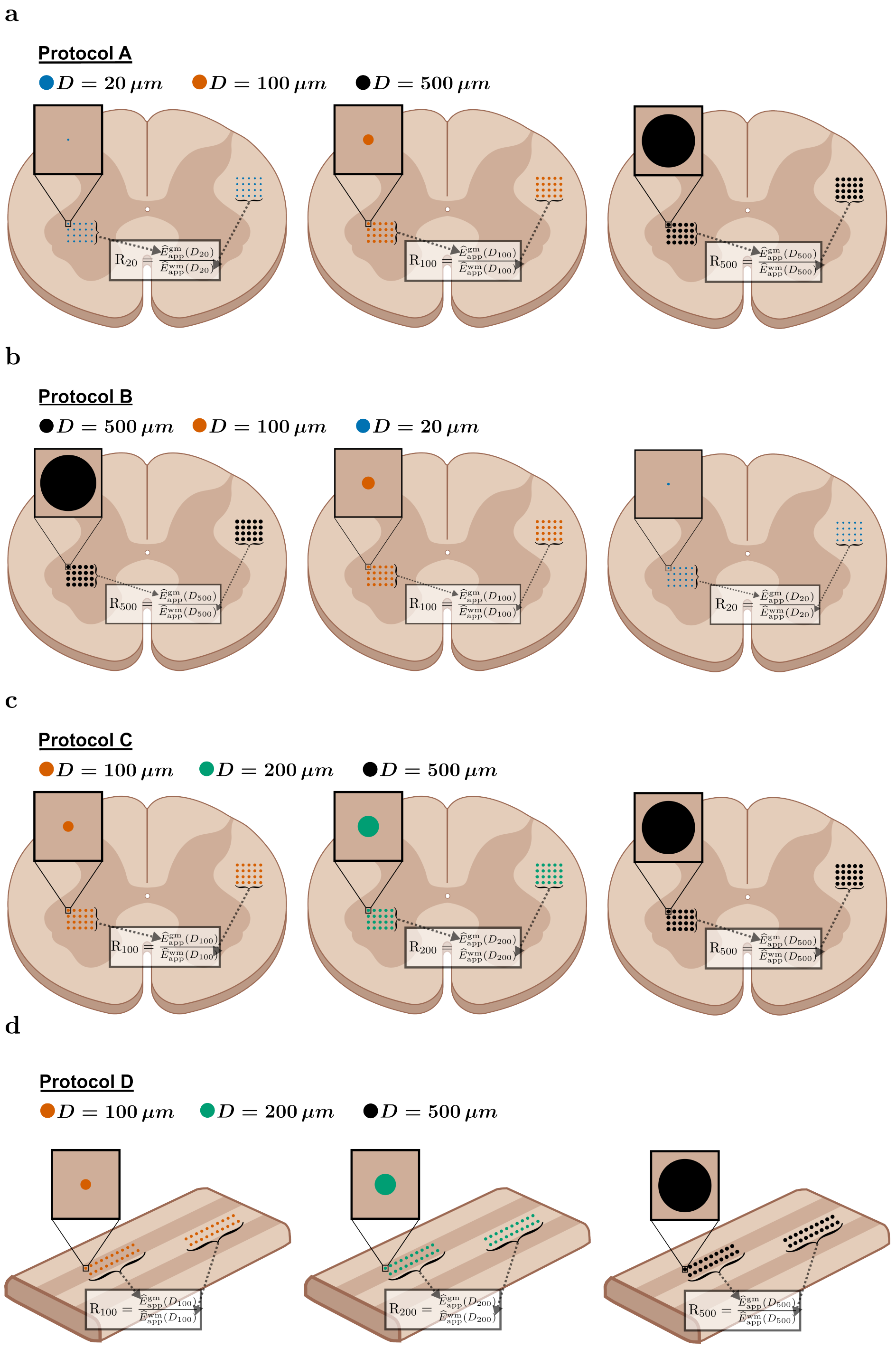


Fig. S3: **Sketches to illustrate the computation of stiffness ratios and the order of measurements for different protocols.** (**a**) Protocol A: D = 20, D = 100, D = 500 $\mu$m, transverse plane. (**b**) Protocol B: D = 500, D = 100, D = 20 $\mu$m, transverse plane. (**c**) Protocol C: D = 100, D = 200, D = 500 $\mu$m, transverse plane. (**d**) Protocol D: D = 100, D = 200, D = 500 $\mu$m, coronal plane. The spinal cord sketches were created using BioRender.

*Validation of experimental approach*

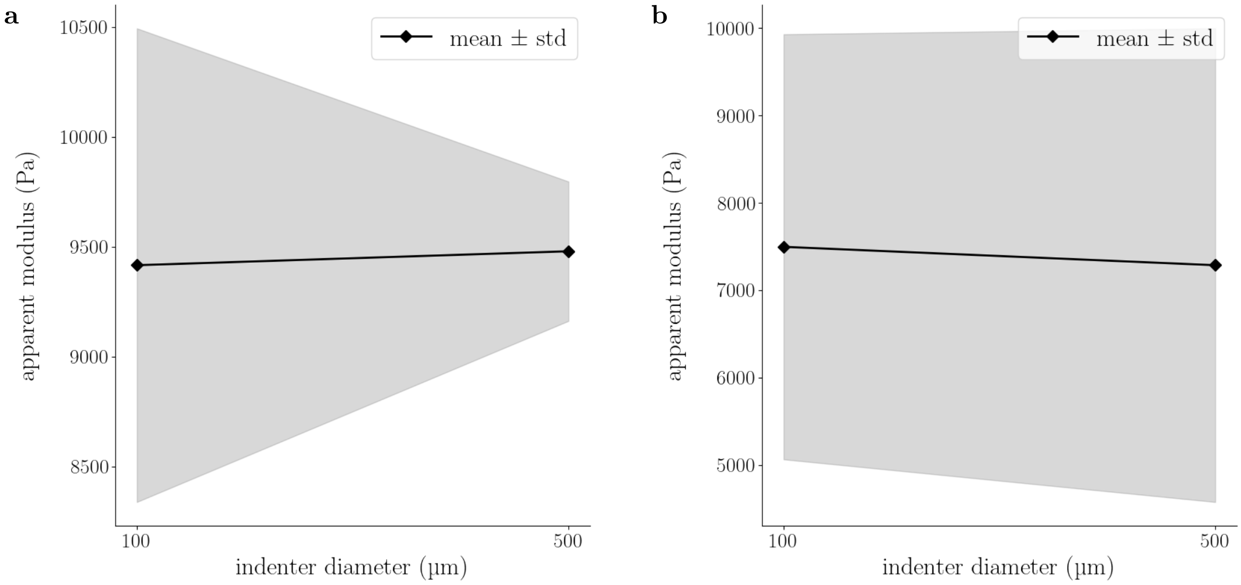


Fig. S4: **Results for tests on agarose with different indenter sizes.** (**a**) Mean values and standard deviations obtained from 20-point indentation grids measured at the same locations on eight 3 % agarose samples, using first a 100 $\mu$m bead and subsequently a 500 $\mu$m bead. (**b**) Mean values and standard deviations obtained from 20-point indentation grids measured at the same locations on eight 3 % agarose samples, using first a 500 $\mu$m bead and subsequently a 100 $\mu$m bead.

*Indenter size-dependence of spinal cord gray and white matter tissue: apparent moduli*

Table S3: **Mean apparent moduli across protocols.**

| **sequence of indenter diameter** | **anatomical plane** | **gray matter stiffness (kPa)** | | | |
|---|---|---|---|---|---|
| | | **D20** | **D100** | **D200** | **D500** |
| **PA**: 20 → 100 → 500 | transverse | 1.57 ± 1.23 | 1.15 ± 0.43 | - | 1.14 ± 0.77 |
| **PB**: 500 → 100 → 20 | transverse | 3.43 ± 3.09 | 1.30 ± 0.72 | - | 0.77 ± 0.30 |
| **PC**: 100 → 200 → 500 | transverse | - | 1.05 ± 0.45 | 0.76 ± 0.33 | 1.07 ± 0.56 |
| **PD**: 100 → 200 → 500 | coronal | - | 0.62 ± 0.16 | 0.53 ± 0.22 | 0.75 ± 0.29 |

| **sequence of indenter diameter** | **anatomical plane** | **white matter stiffness (kPa)** | | | |
|---|---|---|---|---|---|
| | | **D20** | **D100** | **D200** | **D500** |
| **PA**: 20 → 100 → 500 | transverse | 1.73 ± 0.94 | 1.18 ± 0.49 | - | 1.76 ± 1.14 |
| **PB**: 500 → 100 → 20 | transverse | 4.40 ± 4.35 | 1.28 ± 0.65 | - | 1.08 ± 0.54 |
| **PC**: 100 → 200 → 500 | transverse | - | 1.10 ± 0.50 | 0.82 ± 0.50 | 1.62 ± 0.88 |
| **PD**: 100 → 200 → 500 | coronal | - | 0.65 ± 0.24 | 0.56 ± 0.30 | 0.87 ± 0.56 |

Summary of mean apparent moduli and corresponding standard deviations for each test sequence and sample orientation. Magnitudes are highlighted with corresponding colors, with turquoise indicating the lowest and magenta the highest values. The two Protocol B D20 values were not color-coded, since their substantially higher magnitude would impair the readability of the color scale for the remaining entries.

Table S4: **Normalized mean apparent moduli across protocols.**

| sequence of indenter diameter | anatomical plane | normalized gray matter stiffness (-) | | | |
|---|---|---|---|---|---|
| | | D20 | D100 | D200 | D500 |
| **PA**: 20 → 100 → 500 | transverse | 0.91 ± 0.42 | 1.05 ± 0.30 | - | 0.83 ± 0.33 |
| **PB**: 500 → 100 → 20 | transverse | 0.82 ± 0.45 | 1.01 ± 0.41 | - | 0.85 ± 0.30 |
| **PC**: 100 → 200 → 500 | transverse | - | 0.98 ± 0.35 | 0.99 ± 0.40 | 0.79 ± 0.34 |
| **PD**: 100 → 200 → 500 | coronal | - | 0.99 ± 0.24 | 0.99 ± 0.39 | 1.00 ± 0.37 |

| sequence of indenter diameter | anatomical plane | normalized white matter stiffness (-) | | | |
|---|---|---|---|---|---|
| | | D20 | D100 | D200 | D500 |
| **PA**: 20 → 100 → 500 | transverse | 1.05 ± 0.51 | 0.97 ± 0.30 | - | 1.13 ± 0.33 |
| **PB**: 500 → 100 → 20 | transverse | 1.16 ± 0.57 | 0.99 ± 0.42 | - | 1.12 ± 0.39 |
| **PC**: 100 → 200 → 500 | transverse | - | 1.01 ± 0.30 | 1.01 ± 0.43 | 1.15 ± 0.36 |
| **PD**: 100 → 200 → 500 | coronal | - | 1.01 ± 0.37 | 1.01 ± 0.51 | 1.00 ± 0.39 |

Summary of normalized mean apparent moduli and corresponding standard deviations for each test sequence and sample orientation. Colors indicate percentage differences as follows: turquoise, negative values; white, zero; magenta, positive values (shade intensity increases with absolute magnitude)

Table S5: **Changes in mean apparent modulus across protocols.**

| sequence of bead diameter | orientation | gray matter | | | white matter | | |
|---|---|---|---|---|---|---|---|
| | | 20 ⇒ 100 | 100 ⇒ 500 | 20 ⇒ 500 | 20 ⇒ 100 | 100 ⇒ 500 | 20 ⇒ 500 |
| **PA**: 20 → 100 → 500 | transverse | -27% | -1% | -27% | -33% | 52% | 1% |
| **PB**: 500 → 100 → 20 | transverse | -62% | -40% | -77% | -71% | -15% | -75% |
| | | 100 ⇒ 200 | 200 ⇒ 500 | 100 ⇒ 500 | 100 ⇒ 200 | 200 ⇒ 500 | 100 ⇒ 500 |
| **PC**: 100 → 200 → 500 | transverse | -27% | 41% | 2% | -25% | 97% | 47% |
| **PD**: 100 → 200 → 500 | coronal | -15% | 42% | 21% | -15% | 56% | 33% |

Summary of changes in mean apparent moduli for each test sequence and sample orientation. Colors indicate percentage differences as follows: turquoise, negative values; white, zero; magenta, positive values (shade intensity increases with absolute magnitude).

Table S6: **Changes in mean apparent modulus from spinal cord gray to white matter mean tissue**.

| Sequence of indenter diameter | Orientation | Change from gray to white matter stiffness | | | |
|---|---|---|---|---|---|
| | | D20 | D100 | D200 | D500 |
| **PA**: 20 → 100 → 500 | transverse | 10% | 0% | - | 54% |
| **PB**: 500 → 100 → 20 | transverse | 27% | -1% | - | 40% |
| **PC**: 100 → 200 → 500 | transverse | - | 5% | 8% | 51% |
| **PD**: 100 → 200 → 500 | coronal | - | 5% | 5% | 16% |

Summary of changes in mean apparent modulus for each test sequence and sample orientation.

Intensities are highlighted with corresponding colors; turquoise is the lowest and magenta the highest value.

### *Diffusion properties of CNS tissue*

Table S7: **Diffusion properties of CNS tissue**.

| Study | Species | Region | Imaging Modality | GM AD | GM RD | WM AD | WM RD |
|---|---|---|---|---|---|---|---|
| Brander et al. ([64]) | Human | C3 | Axial DTI (3 T) | 1.16 ± 0.14 | 0.68 ± 0.08 | – | – |
| Saksena et al. ([65]) | Human (children) | C3 | Axial DTI (3 T) | 1.24 ± 0.13 | 0.72 ± 0.07 | – | – |
| Al-Shaari et al. ([66]) | Human | C2–C5 | Axial DTI (3 T) | 1.12 ± 0.06 | 0.62 ± 0.05 | – | – |
| Kim et al. ([67]) | Human (*ex vivo*) | Cervical | DTI | 0.47 ± 0.05 | 0.28 ± 0.06 | 0.57 ± 0.04 | 0.17 ± 0.02 |
| Kim et al. ([67]) | Pig (*ex vivo*) | Cervical | DTI | 0.54 ± 0.03 | 0.30 ± 0.02 | 0.83 ± 0.01 | 0.17 ± 0.01 |
| Xu et al. ([68]) | Human | Cervical | Cardiac-gated DTI | 1.11 | 0.55 | 1.74 | – |
| Onu et al. ([69]) | Human | C3–C5 | in-vivo DTI (3 T) | 1.17 | 0.83 | 1.73 | 0.48 |
| Song et al. ([70]) | Mouse (adult) | Corpus callosum WM | MRI DTI (in vivo) | – | – | 1.4 | 0.4 |
| Szczepankiewicz et al. ([71]) | Human (adult) | Brain | MRI DKI (3 T) | 1.0 | 0.80 | 2.0 | 0.31 |
| Lebel & Beaulieu ([72]) | Human (adult) | Brain | MRI DTI (1.5–3 T) | 0.8 | 0.8 | 1.5 | 0.5 |

GM: gray matter, WM: white matter, AD: axial diffusivity, RD: radial diffusivity.

Table S8: **Protocol A: gray matter – overall**

| contrast | diff | p.raw | p.holm |
|---|---|---|---|
| D = 100 $\mu$m / D = 20 $\mu$m | -423 | 0.0202 | 0.1212 |
| D = 500 $\mu$m / D = 20 $\mu$m | -429 | 0.0258 | 0.1291 |
| D = 500 $\mu$m / D = 100 $\mu$m | -5.99 | 0.9460 | 1.0000 |

Table S9: **Protocol A: gray matter – overall, normalized**

| contrast | diff | p.raw | p.holm |
|---|---|---|---|
| D = 100 $\mu$m / D = 20 $\mu$m | 0.136 | 0.0410 | 0.2376 |
| D = 500 $\mu$m / D = 20 $\mu$m | -0.0844 | 0.2108 | 0.4534 |
| D = 500 $\mu$m / D = 100 $\mu$m | -0.22 | 1.5e-06 | 1.2e-05 |

Table S10: **Protocol A: gray matter – sample-wise, 20 $\mu$m**

| sample1 | sample2 | diff | p.adj |
|---|---|---|---|
| 1 | 2 | 3022.8379 | p < 0.0001 |
| 1 | 3 | -58.6334 | 1.0000 |
| 1 | 4 | 28.8817 | 1.0000 |
| 1 | 5 | 439.0947 | 0.9152 |
| 1 | 6 | 136.1675 | 1.0000 |
| 1 | 7 | 46.1287 | 1.0000 |
| 1 | 8 | 219.7668 | 0.9992 |
| 2 | 3 | -3081.4713 | p < 0.0001 |
| 2 | 4 | -2993.9562 | p < 0.0001 |
| 2 | 5 | -2583.7432 | p < 0.0001 |
| 2 | 6 | -2886.6704 | 0.0007 |
| 2 | 7 | -2976.7092 | p < 0.0001 |
| 2 | 8 | -2803.0711 | p < 0.0001 |
| 3 | 4 | 87.5152 | 1.0000 |
| 3 | 5 | 497.7281 | 0.9588 |
| 3 | 6 | 194.8009 | 1.0000 |
| 3 | 7 | 104.7622 | 1.0000 |
| 3 | 8 | 278.4002 | 0.9992 |
| 4 | 5 | 410.2129 | 0.9705 |
| 4 | 6 | 107.2858 | 1.0000 |
| 4 | 7 | 17.247 | 1.0000 |
| 4 | 8 | 190.8851 | 0.9999 |
| 5 | 6 | -302.9272 | 0.9996 |
| 5 | 7 | -392.9659 | 0.9637 |
| 5 | 8 | -219.3279 | 0.9993 |
| 6 | 7 | -90.0388 | 1.0000 |
| 6 | 8 | 83.5993 | 1.0000 |
| 7 | 8 | 173.638 | 0.9999 |

Table S11: **Protocol A: gray matter – sample-wise, 100 $\mu$m**

| sample1 | sample2 | diff | p.adj |
|---|---|---|---|
| 1 | 2 | -187.3634 | 0.8012 |
| 1 | 3 | -496.3826 | 0.0022 |
| 1 | 4 | 16.5551 | 1.0000 |
| 1 | 5 | -198.9909 | 0.6043 |
| 1 | 6 | 668.845 | 0.0038 |
| 1 | 7 | 324.0598 | 0.0973 |
| 1 | 8 | 9.8959 | 1.0000 |
| 2 | 3 | -309.0192 | 0.3214 |
| 2 | 4 | 203.9185 | 0.8382 |
| 2 | 5 | -11.6275 | 1.0000 |
| 2 | 6 | 856.2085 | 0.0002 |
| 2 | 7 | 511.4233 | 0.0038 |
| 2 | 8 | 197.2593 | 0.9075 |
| 3 | 4 | 512.9377 | 0.0088 |
| 3 | 5 | 297.3917 | 0.2397 |
| 3 | 6 | 1165.2277 | $p < 0.0001$ |
| 3 | 7 | 820.4424 | $p < 0.0001$ |
| 3 | 8 | 506.2785 | 0.0275 |
| 4 | 5 | -215.546 | 0.7048 |
| 4 | 6 | 652.2899 | 0.0132 |
| 4 | 7 | 307.5047 | 0.3002 |
| 4 | 8 | -6.6592 | 1.0000 |
| 5 | 6 | 867.836 | 0.0001 |
| 5 | 7 | 523.0507 | 0.0004 |
| 5 | 8 | 208.8868 | 0.8279 |
| 6 | 7 | -344.7852 | 0.5018 |
| 6 | 8 | -658.9492 | 0.0213 |
| 7 | 8 | -314.164 | 0.4019 |

Table S12: **Protocol A: gray matter – sample-wise, 500 $\mu$m**

| sample1 | sample2 | diff | p.adj |
|---|---|---|---|
| 1 | 2 | -518.4944 | 0.1270 |
| 1 | 3 | -537.3576 | 0.0556 |
| 1 | 4 | 9.7733 | 1.0000 |
| 1 | 5 | 245.2566 | 0.8562 |
| 1 | 6 | 878.3298 | 0.0025 |
| 1 | 7 | 1270.3946 | $p < 0.0001$ |
| 1 | 8 | -113.6755 | 0.9997 |
| 2 | 3 | -18.8632 | 1.0000 |
| 2 | 4 | 528.2677 | 0.2037 |
| 2 | 5 | 763.7509 | 0.0034 |
| 2 | 6 | 1396.8242 | $p < 0.0001$ |
| 2 | 7 | 1788.8889 | $p < 0.0001$ |
| 2 | 8 | 404.8189 | 0.7450 |
| 3 | 4 | 547.1309 | 0.1149 |
| 3 | 5 | 782.6141 | 0.0008 |
| 3 | 6 | 1415.6874 | $p < 0.0001$ |
| 3 | 7 | 1807.7522 | $p < 0.0001$ |
| 3 | 8 | 423.6821 | 0.6504 |
| 4 | 5 | 235.4832 | 0.9319 |
| 4 | 6 | 868.5565 | 0.0082 |
| 4 | 7 | 1260.6213 | $p < 0.0001$ |
| 4 | 8 | -123.4488 | 0.9997 |
| 5 | 6 | 633.0733 | 0.0840 |
| 5 | 7 | 1025.138 | $p < 0.0001$ |
| 5 | 8 | -358.9321 | 0.8118 |
| 6 | 7 | 392.0647 | 0.6510 |
| 6 | 8 | -992.0053 | 0.0101 |
| 7 | 8 | -1384.0701 | $p < 0.0001$ |

Table S13: **Protocol A: White matter – overall**

| contrast | diff | p.raw | p.holm |
|---|---|---|---|
| D = 100 $\mu$m / D = 20 $\mu$m | -579 | p < 0.0001 | 0.0001 |
| D = 500 $\mu$m / D = 20 $\mu$m | 24.2 | 0.8778 | 1.0000 |
| D = 500 $\mu$m / D = 100 $\mu$m | 603 | p < 0.0001 | p < 0.0001 |

Table S14: **Protocol A: White matter – overall, normalized**

| contrast | diff | p.raw | p.holm |
|---|---|---|---|
| D = 100 $\mu$m / D = 20 $\mu$m | -0.0838 | 0.1511 | 0.4534 |
| D = 500 $\mu$m / D = 20 $\mu$m | 0.0844 | 0.1583 | 0.4534 |
| D = 500 $\mu$m / D = 100 $\mu$m | 0.168 | 1.6e-05 | 0.0001 |

Table S15: **Protocol A: White matter – sample-wise, 20 $\mu$m**

| sample1 | sample2 | diff | p.adj |
|---|---|---|---|
| 1 | 2 | 2628.6075 | $p < 0.0001$ |
| 1 | 3 | -435.9245 | 0.9531 |
| 1 | 4 | 1050.5717 | 0.2361 |
| 1 | 5 | 354.7053 | 0.9819 |
| 1 | 6 | 622.1556 | 0.7849 |
| 1 | 7 | 152.4218 | 0.9999 |
| 1 | 8 | 678.471 | 0.7019 |
| 2 | 3 | -3064.5321 | $p < 0.0001$ |
| 2 | 4 | -1578.0358 | 0.0049 |
| 2 | 5 | -2273.9023 | $p < 0.0001$ |
| 2 | 6 | -2006.4519 | $p < 0.0001$ |
| 2 | 7 | -2476.1858 | $p < 0.0001$ |
| 2 | 8 | -1950.1365 | $p < 0.0001$ |
| 3 | 4 | 1486.4962 | 0.0041 |
| 3 | 5 | 790.6298 | 0.2216 |
| 3 | 6 | 1058.0801 | 0.0603 |
| 3 | 7 | 588.3463 | 0.6508 |
| 3 | 8 | 1114.3955 | 0.0390 |
| 4 | 5 | -695.8664 | 0.5517 |
| 4 | 6 | -428.4161 | 0.9560 |
| 4 | 7 | -898.1499 | 0.2684 |
| 4 | 8 | -372.1007 | 0.9797 |
| 5 | 6 | 267.4503 | 0.9927 |
| 5 | 7 | -202.2835 | 0.9983 |
| 5 | 8 | 323.7657 | 0.9776 |
| 6 | 7 | -469.7338 | 0.8780 |
| 6 | 8 | 56.3154 | 1.0000 |
| 7 | 8 | 526.0492 | 0.8009 |

Table S16: **Protocol A: White matter – sample-wise, 100 $\mu$m**

| sample1 | sample2 | diff | p.adj |
|---|---|---|---|
| 1 | 2 | -37.7216 | 1.0000 |
| 1 | 3 | -256.1963 | 0.5609 |
| 1 | 4 | 648.5342 | p < 0.0001 |
| 1 | 5 | -356.4017 | 0.1038 |
| 1 | 6 | 621.3883 | p < 0.0001 |
| 1 | 7 | 58.2016 | 0.9997 |
| 1 | 8 | 51.5292 | 0.9999 |
| 2 | 3 | -218.4748 | 0.7324 |
| 2 | 4 | 686.2558 | p < 0.0001 |
| 2 | 5 | -318.6801 | 0.1904 |
| 2 | 6 | 659.1099 | p < 0.0001 |
| 2 | 7 | 95.9232 | 0.9921 |
| 2 | 8 | 89.2508 | 0.9958 |
| 3 | 4 | 904.7305 | p < 0.0001 |
| 3 | 5 | -100.2053 | 0.9965 |
| 3 | 6 | 877.5847 | p < 0.0001 |
| 3 | 7 | 314.398 | 0.2790 |
| 3 | 8 | 307.7256 | 0.3338 |
| 4 | 5 | -1004.9359 | p < 0.0001 |
| 4 | 6 | -27.1458 | 1.0000 |
| 4 | 7 | -590.3326 | 0.0001 |
| 4 | 8 | -597.005 | 0.0001 |
| 5 | 6 | 977.79 | p < 0.0001 |
| 5 | 7 | 414.6033 | 0.0268 |
| 5 | 8 | 407.9309 | 0.0397 |
| 6 | 7 | -563.1867 | 0.0002 |
| 6 | 8 | -569.8591 | 0.0002 |
| 7 | 8 | -6.6724 | 1.0000 |

Table S17: **Protocol A: White matter – sample-wise, 500 $\mu$m**

| sample1 | sample2 | diff | p.adj |
|---|---|---|---|
| 1 | 2 | -490.7332 | 0.1572 |
| 1 | 3 | -257.9992 | 0.9274 |
| 1 | 4 | 423.4263 | 0.3586 |
| 1 | 5 | 369.0916 | 0.5623 |
| 1 | 6 | 2465.3981 | $p < 0.0001$ |
| 1 | 7 | 1758.7288 | $p < 0.0001$ |
| 1 | 8 | 19.0391 | 1.0000 |
| 2 | 3 | 232.734 | 0.9518 |
| 2 | 4 | 914.1594 | 0.0001 |
| 2 | 5 | 859.8248 | 0.0004 |
| 2 | 6 | 2956.1313 | $p < 0.0001$ |
| 2 | 7 | 2249.4619 | $p < 0.0001$ |
| 2 | 8 | 509.7722 | 0.1019 |
| 3 | 4 | 681.4255 | 0.0363 |
| 3 | 5 | 627.0908 | 0.0808 |
| 3 | 6 | 2723.3974 | $p < 0.0001$ |
| 3 | 7 | 2016.728 | $p < 0.0001$ |
| 3 | 8 | 277.0383 | 0.8856 |
| 4 | 5 | -54.3347 | 1.0000 |
| 4 | 6 | 2041.9719 | $p < 0.0001$ |
| 4 | 7 | 1335.3025 | $p < 0.0001$ |
| 4 | 8 | -404.3872 | 0.3815 |
| 5 | 6 | 2096.3066 | $p < 0.0001$ |
| 5 | 7 | 1389.6372 | $p < 0.0001$ |
| 5 | 8 | -350.0525 | 0.5943 |
| 6 | 7 | -706.6694 | 0.0034 |
| 6 | 8 | -2446.3591 | $p < 0.0001$ |
| 7 | 8 | -1739.6897 | $p < 0.0001$ |

Table S18: **Protocol B: gray matter – overall**

| contrast | diff | p.raw | p.holm |
| --- | --- | --- | --- |
| D = 100 $\mu$m / D = 20 $\mu$m | -2.13e+03 | p < 0.0001 | p < 0.0001 |
| D = 500 $\mu$m / D = 20 $\mu$m | -2.66e+03 | p < 0.0001 | p < 0.0001 |
| D = 500 $\mu$m / D = 100 $\mu$m | -524 | p < 0.0001 | p < 0.0001 |

Table S19: **Protocol A: gray matter – overall, normalized**

| contrast | diff | p.raw | p.holm |
| --- | --- | --- | --- |
| D = 100 $\mu$m / D = 20 $\mu$m | 0.19 | 0.0121 | 0.0606 |
| D = 500 $\mu$m / D = 20 $\mu$m | 0.0265 | 0.7054 | 1.0000 |
| D = 500 $\mu$m / D = 100 $\mu$m | -0.164 | 0.0006 | 0.0047 |

Table S20: **Protocol B: gray matter – sample-wise, 20 $\mu$m**

| sample1 | sample2 | diff | p.adj |
|---|---|---|---|
| 9 | 10 | -587.6987 | 0.9996 |
| 9 | 11 | -2861.0263 | 0.6749 |
| 9 | 12 | -3412.2142 | 0.7826 |
| 9 | 13 | -1673.3108 | 0.9176 |
| 9 | 14 | -3454.6547 | 0.5908 |
| 9 | 15 | -4061.3259 | 0.1650 |
| 9 | 16 | -3566.1521 | 0.4053 |
| 10 | 11 | -2273.3276 | 0.8834 |
| 10 | 12 | -2824.5155 | 0.9109 |
| 10 | 13 | -1085.6121 | 0.9940 |
| 10 | 14 | -2866.956 | 0.8031 |
| 10 | 15 | -3473.6272 | 0.3755 |
| 10 | 16 | -2978.4534 | 0.6631 |
| 11 | 12 | -551.1879 | 1.0000 |
| 11 | 13 | 1187.7155 | 0.9979 |
| 11 | 14 | -593.6284 | 1.0000 |
| 11 | 15 | -1200.2995 | 0.9986 |
| 11 | 16 | -705.1258 | 1.0000 |
| 12 | 13 | 1738.9034 | 0.9952 |
| 12 | 14 | -42.4405 | 1.0000 |
| 12 | 15 | -649.1116 | 1.0000 |
| 12 | 16 | -153.9379 | 1.0000 |
| 13 | 14 | -1781.3439 | 0.9864 |
| 13 | 15 | -2388.015 | 0.8568 |
| 13 | 16 | -1892.8413 | 0.9672 |
| 14 | 15 | -606.6711 | 1.0000 |
| 14 | 16 | -111.4974 | 1.0000 |
| 15 | 16 | 495.1738 | 1.0000 |

Table S21: **Protocol B: gray matter – sample-wise, 100 $\mu$m**

| sample1 | sample2 | diff | p.adj |
|---|---|---|---|
| 9 | 10 | -298.6988 | 0.7739 |
| 9 | 11 | -673.364 | 0.0142 |
| 9 | 12 | -542.7028 | 0.2235 |
| 9 | 13 | -469.3519 | 0.3187 |
| 9 | 14 | 708.1272 | 0.0153 |
| 9 | 15 | -250.2992 | 0.9272 |
| 9 | 16 | -655.6474 | 0.1667 |
| 10 | 11 | -374.6652 | 0.4999 |
| 10 | 12 | -244.004 | 0.9520 |
| 10 | 13 | -170.6531 | 0.9907 |
| 10 | 14 | 1006.8259 | 0.0001 |
| 10 | 15 | 48.3996 | 1.0000 |
| 10 | 16 | -356.9486 | 0.8435 |
| 11 | 12 | 130.6612 | 0.9988 |
| 11 | 13 | 204.0121 | 0.9739 |
| 11 | 14 | 1381.4911 | $p < 0.0001$ |
| 11 | 15 | 423.0648 | 0.4405 |
| 11 | 16 | 17.7166 | 1.0000 |
| 12 | 13 | 73.3509 | 1.0000 |
| 12 | 14 | 1250.8299 | $p < 0.0001$ |
| 12 | 15 | 292.4036 | 0.9109 |
| 12 | 16 | -112.9446 | 0.9999 |
| 13 | 14 | 1177.479 | $p < 0.0001$ |
| 13 | 15 | 219.0527 | 0.9732 |
| 13 | 16 | -186.2955 | 0.9965 |
| 14 | 15 | -958.4264 | 0.0005 |
| 14 | 16 | -1363.7746 | $p < 0.0001$ |
| 15 | 16 | -405.3482 | 0.7801 |

Table S22: **Protocol B: gray matter – sample-wise, 500 $\mu$m**

| sample1 | sample2 | diff | p.adj |
|---|---|---|---|
| 9 | 10 | 41.7268 | 0.9988 |
| 9 | 11 | -226.8539 | 0.0294 |
| 9 | 12 | 522.9334 | $p < 0.0001$ |
| 9 | 13 | 65.1554 | 0.9882 |
| 9 | 14 | 146.8913 | 0.4659 |
| 9 | 15 | -285.1043 | 0.0073 |
| 9 | 16 | 10.5682 | 1.0000 |
| 10 | 11 | -268.5806 | 0.0036 |
| 10 | 12 | 481.2066 | $p < 0.0001$ |
| 10 | 13 | 23.4287 | 1.0000 |
| 10 | 14 | 105.1645 | 0.8202 |
| 10 | 15 | -326.831 | 0.0009 |
| 10 | 16 | -31.1585 | 1.0000 |
| 11 | 12 | 749.7872 | $p < 0.0001$ |
| 11 | 13 | 292.0093 | 0.0035 |
| 11 | 14 | 373.7451 | $p < 0.0001$ |
| 11 | 15 | -58.2504 | 0.9943 |
| 11 | 16 | 237.4221 | 0.1409 |
| 12 | 13 | -457.7779 | $p < 0.0001$ |
| 12 | 14 | -376.0421 | 0.0002 |
| 12 | 15 | -808.0376 | $p < 0.0001$ |
| 12 | 16 | -512.3651 | $p < 0.0001$ |
| 13 | 14 | 81.7358 | 0.9634 |
| 13 | 15 | -350.2597 | 0.0009 |
| 13 | 16 | -54.5872 | 0.9990 |
| 14 | 15 | -431.9955 | $p < 0.0001$ |
| 14 | 16 | -136.3231 | 0.8081 |
| 15 | 16 | 295.6725 | 0.0452 |

Table S23: **Protocol B: White matter – overall**

| contrast | diff | p.raw | p.holm |
|---|---|---|---|
| D = 100 $\mu$m / D = 20 $\mu$m | -3.08e+03 | p < 0.0001 | p < 0.0001 |
| D = 500 $\mu$m / D = 20 $\mu$m | -3.27e+03 | p < 0.0001 | p < 0.0001 |
| D = 500 $\mu$m / D = 100 $\mu$m | -197 | 0.0065 | 0.0196 |

Table S24: **Protocol A: White matter – overall, normalized**

| contrast | diff | p.raw | p.holm |
|---|---|---|---|
| D = 100 $\mu$m / D = 20 $\mu$m | -0.17 | 0.0492 | 0.1968 |
| D = 500 $\mu$m / D = 20 $\mu$m | -0.0328 | 0.6967 | 1.0000 |
| D = 500 $\mu$m / D = 100 $\mu$m | 0.137 | 0.0047 | 0.0280 |

Table S25: **Protocol B: White matter – sample-wise, 20 $\mu$m**

| sample1 | sample2 | diff | p.adj |
|---|---|---|---|
| 9 | 10 | -3140.1055 | 0.4029 |
| 9 | 11 | -6230.9973 | 0.0125 |
| 9 | 12 | -8831.5123 | 0.1394 |
| 9 | 13 | -7435.7091 | 0.0015 |
| 9 | 14 | -8163.5753 | $p < 0.0001$ |
| 9 | 15 | -7857.2826 | 0.0001 |
| 9 | 16 | -9140.7986 | $p < 0.0001$ |
| 10 | 11 | -3090.8918 | 0.6517 |
| 10 | 12 | -5691.4068 | 0.6633 |
| 10 | 13 | -4295.6036 | 0.2462 |
| 10 | 14 | -5023.4698 | 0.0209 |
| 10 | 15 | -4717.1771 | 0.0660 |
| 10 | 16 | -6000.6931 | 0.0070 |
| 11 | 12 | -2600.515 | 0.9940 |
| 11 | 13 | -1204.7118 | 0.9985 |
| 11 | 14 | -1932.578 | 0.9391 |
| 11 | 15 | -1626.2852 | 0.9822 |
| 11 | 16 | -2909.8013 | 0.7162 |
| 12 | 13 | 1395.8032 | 0.9999 |
| 12 | 14 | 667.937 | 1.0000 |
| 12 | 15 | 974.2297 | 1.0000 |
| 12 | 16 | -309.2863 | 1.0000 |
| 13 | 14 | -727.8662 | 0.9998 |
| 13 | 15 | -421.5734 | 1.0000 |
| 13 | 16 | -1705.0895 | 0.9767 |
| 14 | 15 | 306.2928 | 1.0000 |
| 14 | 16 | -977.2233 | 0.9971 |
| 15 | 16 | -1283.516 | 0.9904 |

Table S26: **Protocol B: White matter – sample-wise, 100 $\mu$m**

| sample1 | sample2 | diff | p.adj |
|---|---|---|---|
| 9 | 10 | -174.6965 | 0.9786 |
| 9 | 11 | -654.1757 | 0.0073 |
| 9 | 12 | -804.8327 | 0.0160 |
| 9 | 13 | -1024.3637 | p < 0.0001 |
| 9 | 14 | -362.3746 | 0.4505 |
| 9 | 15 | 190.6928 | 0.9651 |
| 9 | 16 | -482.5895 | 0.1217 |
| 10 | 11 | -479.4793 | 0.1385 |
| 10 | 12 | -630.1362 | 0.1344 |
| 10 | 13 | -849.6672 | 0.0008 |
| 10 | 14 | -187.6782 | 0.9658 |
| 10 | 15 | 365.3893 | 0.4925 |
| 10 | 16 | -307.893 | 0.6739 |
| 11 | 12 | -150.657 | 0.9980 |
| 11 | 13 | -370.188 | 0.5340 |
| 11 | 14 | 291.8011 | 0.7031 |
| 11 | 15 | 844.8685 | 0.0002 |
| 11 | 16 | 171.5862 | 0.9758 |
| 12 | 13 | -219.531 | 0.9857 |
| 12 | 14 | 442.4581 | 0.5397 |
| 12 | 15 | 995.5255 | 0.0010 |
| 12 | 16 | 322.2432 | 0.8564 |
| 13 | 14 | 661.9891 | 0.0169 |
| 13 | 15 | 1215.0565 | p < 0.0001 |
| 13 | 16 | 541.7742 | 0.0984 |
| 14 | 15 | 553.0674 | 0.0487 |
| 14 | 16 | -120.2148 | 0.9971 |
| 15 | 16 | -673.2823 | 0.0060 |

Table S27: **Protocol B: White matter – sample-wise, 500 $\mu$m**

| sample1 | sample2 | diff | p.adj |
|---|---|---|---|
| 9 | 10 | 119.8091 | 0.9865 |
| 9 | 11 | -852.4281 | $p < 0.0001$ |
| 9 | 12 | -176.6924 | 0.8428 |
| 9 | 13 | -1103.4271 | $p < 0.0001$ |
| 9 | 14 | -689.3846 | $p < 0.0001$ |
| 9 | 15 | -285.2362 | 0.3359 |
| 9 | 16 | -673.7657 | $p < 0.0001$ |
| 10 | 11 | -972.2372 | $p < 0.0001$ |
| 10 | 12 | -296.5015 | 0.3709 |
| 10 | 13 | -1223.2362 | $p < 0.0001$ |
| 10 | 14 | -809.1937 | $p < 0.0001$ |
| 10 | 15 | -405.0453 | 0.0795 |
| 10 | 16 | -793.5748 | $p < 0.0001$ |
| 11 | 12 | 675.7357 | $p < 0.0001$ |
| 11 | 13 | -250.9989 | 0.5517 |
| 11 | 14 | 163.0435 | 0.8839 |
| 11 | 15 | 567.1919 | 0.0005 |
| 11 | 16 | 178.6624 | 0.8448 |
| 12 | 13 | -926.7346 | $p < 0.0001$ |
| 12 | 14 | -512.6922 | 0.0015 |
| 12 | 15 | -108.5438 | 0.9903 |
| 12 | 16 | -497.0733 | 0.0035 |
| 13 | 14 | 414.0424 | 0.0424 |
| 13 | 15 | 818.1908 | $p < 0.0001$ |
| 13 | 16 | 429.6613 | 0.0374 |
| 14 | 15 | 404.1484 | 0.0385 |
| 14 | 16 | 15.6189 | 1.0000 |
| 15 | 16 | -388.5295 | 0.0665 |

Table S28: **Protocol C: gray matter – overall**

| contrast | diff | p.raw | p.holm |
|---|---|---|---|
| D = 100 $\mu$m / D = 20 $\mu$m | -288 | p < 0.0001 | p < 0.0001 |
| D = 500 $\mu$m / D = 20 $\mu$m | 21.8 | 0.7530 | 0.7715 |
| D = 500 $\mu$m / D = 100 $\mu$m | 310 | p < 0.0001 | p < 0.0001 |

Table S29: **Protocol C: gray matter – overall, normalized**

| contrast | diff | p.raw | p.holm |
|---|---|---|---|
| D = 100 $\mu$m / D = 20 $\mu$m | 0.00836 | 0.8687 | 1.0000 |
| D = 500 $\mu$m / D = 20 $\mu$m | -0.185 | 0.0001 | 0.0009 |
| D = 500 $\mu$m / D = 100 $\mu$m | -0.193 | 0.0001 | 0.0010 |

Table S30: **Protocol C: gray matter – sample-wise, 100 $\mu$m**

| sample1 | sample2 | diff | p.adj |
|---|---|---:|---:|
| 17 | 18 | 176.7771 | 0.8702 |
| 17 | 19 | 295.9428 | 0.2850 |
| 17 | 20 | 848.5025 | $p < 0.0001$ |
| 17 | 21 | -25.6918 | 1.0000 |
| 17 | 22 | 195.977 | 0.9657 |
| 17 | 23 | 128.483 | 0.9712 |
| 17 | 24 | -270.0247 | 0.4844 |
| 18 | 19 | 119.1657 | 0.9684 |
| 18 | 20 | 671.7254 | 0.0001 |
| 18 | 21 | -202.4688 | 0.9687 |
| 18 | 22 | 19.1999 | 1.0000 |
| 18 | 23 | -48.2941 | 0.9999 |
| 18 | 24 | -446.8018 | 0.0116 |
| 19 | 20 | 552.5597 | 0.0018 |
| 19 | 21 | -321.6345 | 0.7178 |
| 19 | 22 | -99.9658 | 0.9992 |
| 19 | 23 | -167.4598 | 0.8113 |
| 19 | 24 | -565.9675 | 0.0003 |
| 20 | 21 | -874.1942 | 0.0014 |
| 20 | 22 | -652.5255 | 0.0209 |
| 20 | 23 | -720.0195 | $p < 0.0001$ |
| 20 | 24 | -1118.5272 | $p < 0.0001$ |
| 21 | 22 | 221.6688 | 0.9824 |
| 21 | 23 | 154.1748 | 0.9932 |
| 21 | 24 | -244.333 | 0.9237 |
| 22 | 23 | -67.494 | 0.9999 |
| 22 | 24 | -466.0017 | 0.1922 |
| 23 | 24 | -398.5077 | 0.0299 |

Table S31: **Protocol C: gray matter – sample-wise, 200 $\mu$m**

| sample1 | sample2 | diff | p.adj |
|---|---|---|---|
| 17 | 18 | 33.8463 | 1.0000 |
| 17 | 19 | 501.1194 | $p < 0.0001$ |
| 17 | 20 | 103.7831 | 0.9790 |
| 17 | 21 | 49.1829 | 1.0000 |
| 17 | 22 | 373.8758 | 0.1056 |
| 17 | 23 | -53.3504 | 0.9993 |
| 17 | 24 | -96.9881 | 0.9805 |
| 18 | 19 | 467.2731 | $p < 0.0001$ |
| 18 | 20 | 69.9368 | 0.9968 |
| 18 | 21 | 15.3366 | 1.0000 |
| 18 | 22 | 340.0295 | 0.1385 |
| 18 | 23 | -87.1967 | 0.9724 |
| 18 | 24 | -130.8344 | 0.8544 |
| 19 | 20 | -397.3364 | 0.0031 |
| 19 | 21 | -451.9365 | 0.0123 |
| 19 | 22 | -127.2436 | 0.9726 |
| 19 | 23 | -554.4699 | $p < 0.0001$ |
| 19 | 24 | -598.1075 | $p < 0.0001$ |
| 20 | 21 | -54.6002 | 0.9999 |
| 20 | 22 | 270.0927 | 0.4876 |
| 20 | 23 | -157.1335 | 0.7537 |
| 20 | 24 | -200.7712 | 0.5417 |
| 21 | 22 | 324.6929 | 0.4340 |
| 21 | 23 | -102.5333 | 0.9919 |
| 21 | 24 | -146.171 | 0.9510 |
| 22 | 23 | -427.2262 | 0.0210 |
| 22 | 24 | -470.8639 | 0.0109 |
| 23 | 24 | -43.6377 | 0.9998 |

Table S32: **Protocol C: gray matter – sample-wise, 500 $\mu$m**

| sample1 | sample2 | diff | p.adj |
|---|---|---|---|
| 17 | 18 | 318.8452 | 0.5180 |
| 17 | 19 | 757.2157 | 0.0003 |
| 17 | 20 | 98.2486 | 0.9993 |
| 17 | 21 | 56.3349 | 1.0000 |
| 17 | 22 | 368.1688 | 0.7321 |
| 17 | 23 | -184.3967 | 0.9514 |
| 17 | 24 | -275.0125 | 0.8059 |
| 18 | 19 | 438.3705 | 0.0602 |
| 18 | 20 | -220.5965 | 0.8764 |
| 18 | 21 | -262.5102 | 0.9207 |
| 18 | 22 | 49.3236 | 1.0000 |
| 18 | 23 | -503.2418 | 0.0191 |
| 18 | 24 | -593.8576 | 0.0131 |
| 19 | 20 | -658.9671 | 0.0026 |
| 19 | 21 | -700.8807 | 0.0292 |
| 19 | 22 | -389.0469 | 0.6051 |
| 19 | 23 | -941.6124 | $p < 0.0001$ |
| 19 | 24 | -1032.2282 | $p < 0.0001$ |
| 20 | 21 | -41.9137 | 1.0000 |
| 20 | 22 | 269.9201 | 0.9319 |
| 20 | 23 | -282.6453 | 0.6781 |
| 20 | 24 | -373.2611 | 0.4637 |
| 21 | 22 | 311.8338 | 0.9361 |
| 21 | 23 | -240.7316 | 0.9505 |
| 21 | 24 | -331.3474 | 0.8331 |
| 22 | 23 | -552.5654 | 0.1771 |
| 22 | 24 | -643.1812 | 0.1045 |
| 23 | 24 | -90.6158 | 0.9994 |

Table S33: **Protocol C: White matter – overall**

| contrast | diff | p.raw | p.holm |
|---|---|---|---|
| D = 100 $\mu$m / D = 20 $\mu$m | -281 | p < 0.0001 | p < 0.0001 |
| D = 500 $\mu$m / D = 20 $\mu$m | 515 | p < 0.0001 | p < 0.0001 |
| D = 500 $\mu$m / D = 100 $\mu$m | 797 | p < 0.0001 | p < 0.0001 |

Table S34: **Protocol C: White matter – overall, normalized**

| contrast | diff | p.raw | p.holm |
|---|---|---|---|
| D = 100 $\mu$m / D = 20 $\mu$m | -0.00507 | 0.9047 | 1.0000 |
| D = 500 $\mu$m / D = 20 $\mu$m | 0.139 | 0.0004 | 0.0022 |
| D = 500 $\mu$m / D = 100 $\mu$m | 0.144 | 0.0020 | 0.0098 |

Table S35: **Protocol C: White matter – sample-wise, 100 $\mu$m**

| sample1 | sample2 | diff | p.adj |
|---|---|---|---|
| 17 | 18 | 892.04 | $p < 0.0001$ |
| 17 | 19 | 122.1363 | 0.9327 |
| 17 | 20 | 88.023 | 0.9907 |
| 17 | 21 | -229.223 | 0.3515 |
| 17 | 22 | 533.1816 | $p < 0.0001$ |
| 17 | 23 | 451.2022 | 0.0005 |
| 17 | 24 | -346.6829 | 0.0199 |
| 18 | 19 | -769.9037 | $p < 0.0001$ |
| 18 | 20 | -804.017 | $p < 0.0001$ |
| 18 | 21 | -1121.263 | $p < 0.0001$ |
| 18 | 22 | -358.8584 | 0.0136 |
| 18 | 23 | -440.8378 | 0.0008 |
| 18 | 24 | -1238.7229 | $p < 0.0001$ |
| 19 | 20 | -34.1133 | 1.0000 |
| 19 | 21 | -351.3593 | 0.0198 |
| 19 | 22 | 411.0453 | 0.0023 |
| 19 | 23 | 329.0659 | 0.0335 |
| 19 | 24 | -468.8192 | 0.0003 |
| 20 | 21 | -317.246 | 0.0647 |
| 20 | 22 | 445.1587 | 0.0010 |
| 20 | 23 | 363.1792 | 0.0161 |
| 20 | 24 | -434.7059 | 0.0015 |
| 21 | 22 | 762.4046 | $p < 0.0001$ |
| 21 | 23 | 680.4252 | $p < 0.0001$ |
| 21 | 24 | -117.4599 | 0.9485 |
| 22 | 23 | -81.9794 | 0.9928 |
| 22 | 24 | -879.8645 | $p < 0.0001$ |
| 23 | 24 | -797.8851 | $p < 0.0001$ |

Table S36: **Protocol C: White matter – sample-wise, 200 $\mu$m**

| sample1 | sample2 | diff | p.adj |
|---|---|---:|---:|
| 17 | 18 | 935.0077 | $p < 0.0001$ |
| 17 | 19 | 537.8044 | 0.0003 |
| 17 | 20 | 176.5702 | 0.8345 |
| 17 | 21 | 3.9068 | 1.0000 |
| 17 | 22 | 778.7428 | $p < 0.0001$ |
| 17 | 23 | 282.9089 | 0.2604 |
| 17 | 24 | 43.4304 | 1.0000 |
| 18 | 19 | -397.2034 | 0.0232 |
| 18 | 20 | -758.4375 | $p < 0.0001$ |
| 18 | 21 | -931.1009 | $p < 0.0001$ |
| 18 | 22 | -156.2649 | 0.8925 |
| 18 | 23 | -652.0988 | $p < 0.0001$ |
| 18 | 24 | -891.5774 | $p < 0.0001$ |
| 19 | 20 | -361.2341 | 0.0626 |
| 19 | 21 | -533.8976 | 0.0003 |
| 19 | 22 | 240.9385 | 0.4515 |
| 19 | 23 | -254.8955 | 0.3760 |
| 19 | 24 | -494.374 | 0.0022 |
| 20 | 21 | -172.6634 | 0.8415 |
| 20 | 22 | 602.1726 | $p < 0.0001$ |
| 20 | 23 | 106.3387 | 0.9873 |
| 20 | 24 | -133.1399 | 0.9639 |
| 21 | 22 | 774.836 | $p < 0.0001$ |
| 21 | 23 | 279.0021 | 0.2615 |
| 21 | 24 | 39.5236 | 1.0000 |
| 22 | 23 | -495.8339 | 0.0011 |
| 22 | 24 | -735.3125 | $p < 0.0001$ |
| 23 | 24 | -239.4785 | 0.5162 |

Table S37: **Protocol C: White matter – sample-wise, 500 $\mu$m**

| sample1 | sample2 | diff | p.adj |
|---|---|---|---|
| 17 | 18 | 2002.506 | p < 0.0001 |
| 17 | 19 | 1245.8014 | p < 0.0001 |
| 17 | 20 | 384.223 | 0.1962 |
| 17 | 21 | 64.407 | 0.9999 |
| 17 | 22 | 1729.9621 | p < 0.0001 |
| 17 | 23 | 92.6433 | 0.9986 |
| 17 | 24 | 561.7584 | 0.0133 |
| 18 | 19 | -756.7046 | 0.0001 |
| 18 | 20 | -1618.2831 | p < 0.0001 |
| 18 | 21 | -1938.0991 | p < 0.0001 |
| 18 | 22 | -272.5439 | 0.6778 |
| 18 | 23 | -1909.8628 | p < 0.0001 |
| 18 | 24 | -1440.7477 | p < 0.0001 |
| 19 | 20 | -861.5784 | p < 0.0001 |
| 19 | 21 | -1181.3944 | p < 0.0001 |
| 19 | 22 | 484.1607 | 0.0679 |
| 19 | 23 | -1153.1581 | p < 0.0001 |
| 19 | 24 | -684.0431 | 0.0016 |
| 20 | 21 | -319.816 | 0.4392 |
| 20 | 22 | 1345.7391 | p < 0.0001 |
| 20 | 23 | -291.5797 | 0.5454 |
| 20 | 24 | 177.5354 | 0.9561 |
| 21 | 22 | 1665.5552 | p < 0.0001 |
| 21 | 23 | 28.2363 | 1.0000 |
| 21 | 24 | 497.3514 | 0.0499 |
| 22 | 23 | -1637.3188 | p < 0.0001 |
| 22 | 24 | -1168.2038 | p < 0.0001 |
| 23 | 24 | 469.1151 | 0.0729 |

Table S38: **Protocol D: gray matter – overall**

| contrast | diff | p.raw | p.holm |
|---|---|---|---|
| D = 200 $\mu$m vs D = 100 $\mu$m | -91.5 | 0.0002 | 0.0008 |
| D = 500 $\mu$m vs D = 100 $\mu$m | 129 | <0.0001 | <0.0001 |
| D = 500 $\mu$m vs D = 200 $\mu$m | 221 | <0.0001 | <0.0001 |

Table S39: **Protocol D: gray matter – overall, normalized**

| contrast | diff | p.raw | p.holm |
|---|---|---|---|
| D = 100 $\mu$m / D = 20 $\mu$m | -0.000248 | 0.9950 | 1.0000 |
| D = 500 $\mu$m / D = 20 $\mu$m | 0.00122 | 0.9735 | 1.0000 |
| D = 500 $\mu$m / D = 100 $\mu$m | 0.00146 | 0.9739 | 1.0000 |

Table S40: **Protocol D: gray matter – sample-wise, 100 $\mu$m**

| sample1 | sample2 | diff | p.adj |
|---|---|---|---|
| 25 | 26 | 121.3437 | 0.3290 |
| 25 | 27 | 283.0185 | <0.0001 |
| 25 | 28 | 10.8896 | 1.0000 |
| 25 | 29 | -1.4504 | 1.0000 |
| 25 | 30 | 116.0515 | 0.2354 |
| 25 | 31 | 65.1818 | 0.8783 |
| 25 | 32 | -13.7256 | 1.0000 |
| 26 | 27 | 161.6748 | 0.0980 |
| 26 | 28 | -110.4541 | 0.4105 |
| 26 | 29 | -122.7941 | 0.2993 |
| 26 | 30 | -5.2922 | 1.0000 |
| 26 | 31 | -56.1619 | 0.9648 |
| 26 | 32 | -135.0693 | 0.1905 |
| 27 | 28 | -272.1289 | <0.0001 |
| 27 | 29 | -284.469 | <0.0001 |
| 27 | 30 | -166.9671 | 0.0315 |
| 27 | 31 | -217.8367 | 0.0013 |
| 27 | 32 | -296.7441 | <0.0001 |
| 28 | 29 | -12.3401 | 1.0000 |
| 28 | 30 | 105.1618 | 0.3039 |
| 28 | 31 | 54.2921 | 0.9392 |
| 28 | 32 | -24.6152 | 0.9995 |
| 29 | 30 | 117.5019 | 0.2055 |
| 29 | 31 | 66.6322 | 0.8565 |
| 29 | 32 | -12.2752 | 1.0000 |
| 30 | 31 | -50.8697 | 0.9595 |
| 30 | 32 | -129.7771 | 0.1148 |
| 31 | 32 | -78.9074 | 0.7153 |

Table S41: **Protocol D: gray matter – sample-wise, 200 $\mu$m**

| sample1 | sample2 | diff | p.adj |
|---|---|---:|---:|
| 25 | 26 | 117.6126 | 0.7418 |
| 25 | 27 | 209.0778 | 0.0388 |
| 25 | 28 | 185.6567 | 0.0916 |
| 25 | 29 | 196.5239 | 0.0594 |
| 25 | 30 | -68.4381 | 0.9945 |
| 25 | 31 | 321.5629 | 0.0001 |
| 25 | 32 | 92.9444 | 0.8510 |
| 26 | 27 | 91.4652 | 0.9088 |
| 26 | 28 | 68.0442 | 0.9799 |
| 26 | 29 | 78.9114 | 0.9546 |
| 26 | 30 | -186.0506 | 0.4989 |
| 26 | 31 | 203.9504 | 0.0972 |
| 26 | 32 | -24.6681 | 1.0000 |
| 27 | 28 | -23.4211 | 1.0000 |
| 27 | 29 | -12.5539 | 1.0000 |
| 27 | 30 | -277.5159 | 0.0434 |
| 27 | 31 | 112.4851 | 0.6675 |
| 27 | 32 | -116.1334 | 0.6303 |
| 28 | 29 | 10.8672 | 1.0000 |
| 28 | 30 | -254.0948 | 0.0837 |
| 28 | 31 | 135.9062 | 0.4096 |
| 28 | 32 | -92.7123 | 0.8347 |
| 29 | 30 | -264.962 | 0.0606 |
| 29 | 31 | 125.039 | 0.5209 |
| 29 | 32 | -103.5795 | 0.7412 |
| 30 | 31 | 390.001 | 0.0006 |
| 30 | 32 | 161.3825 | 0.6066 |
| 31 | 32 | -228.6185 | 0.0137 |

Table S42: **Protocol D: gray matter – sample-wise, 500 $\mu$m**

| sample1 | sample2 | diff | p.adj |
|---|---|---:|---:|
| 25 | 26 | 193.6161 | 0.2218 |
| 25 | 27 | 242.3304 | 0.0497 |
| 25 | 28 | 230.899 | 0.0736 |
| 25 | 29 | 133.9029 | 0.6727 |
| 25 | 30 | 587.6304 | <0.0001 |
| 25 | 31 | 192.8026 | 0.2122 |
| 25 | 32 | 48.6956 | 0.9986 |
| 26 | 27 | 48.7143 | 0.9987 |
| 26 | 28 | 37.283 | 0.9998 |
| 26 | 29 | -59.7131 | 0.9949 |
| 26 | 30 | 394.0144 | <0.0001 |
| 26 | 31 | -0.8135 | 1.0000 |
| 26 | 32 | -144.9205 | 0.6096 |
| 27 | 28 | -11.4313 | 1.0000 |
| 27 | 29 | -108.4274 | 0.8664 |
| 27 | 30 | 345.3001 | 0.0006 |
| 27 | 31 | -49.5278 | 0.9984 |
| 27 | 32 | -193.6348 | 0.2359 |
| 28 | 29 | -96.9961 | 0.9212 |
| 28 | 30 | 356.7314 | 0.0003 |
| 28 | 31 | -38.0964 | 0.9997 |
| 28 | 32 | -182.2034 | 0.3092 |
| 29 | 30 | 453.7275 | <0.0001 |
| 29 | 31 | 58.8997 | 0.9949 |
| 29 | 32 | -85.2073 | 0.9597 |
| 30 | 31 | -394.8278 | <0.0001 |
| 30 | 32 | -538.9348 | <0.0001 |
| 31 | 32 | -144.107 | 0.6009 |

Table S43: **Protocol D: White matter – overall**

| contrast | diff | p.raw | p.holm |
|---|---|---|---|
| D = 200 $\mu$m vs D = 100 $\mu$m | -96.3 | 0.0072 | 0.0288 |
| D = 500 $\mu$m vs D = 100 $\mu$m | 216 | <0.0001 | 0.0006 |
| D = 500 $\mu$m vs D = 200 $\mu$m | 312 | <0.0001 | <0.0001 |

Table S44: **Protocol D: White matter – overall, normalized**

| contrast | diff | p.raw | p.holm |
|---|---|---|---|
| D = 100 $\mu$m / D = 20 $\mu$m | -0.000113 | 0.9983 | 1.0000 |
| D = 500 $\mu$m / D = 20 $\mu$m | -0.000821 | 0.9859 | 1.0000 |
| D = 500 $\mu$m / D = 100 $\mu$m | -0.000707 | 0.9897 | 1.0000 |

Table S45: **Protocol D: White matter – sample-wise, 100 $\mu$m**

| sample1 | sample2 | diff | p.adj |
|---|---|---|---|
| 25 | 26 | 132.2159 | 0.7961 |
| 25 | 27 | 27.1416 | 1.0000 |
| 25 | 28 | -295.807 | 0.0181 |
| 25 | 29 | -99.5214 | 0.9615 |
| 25 | 30 | -134.2859 | 0.7829 |
| 25 | 31 | -242.6296 | 0.1080 |
| 25 | 32 | -345.2451 | 0.0059 |
| 26 | 27 | -105.0742 | 0.9285 |
| 26 | 28 | -428.0229 | <0.0001 |
| 26 | 29 | -231.7373 | 0.1081 |
| 26 | 30 | -266.5017 | 0.0156 |
| 26 | 31 | -374.8454 | 0.0001 |
| 26 | 32 | -477.4609 | <0.0001 |
| 27 | 28 | -322.9487 | 0.0065 |
| 27 | 29 | -126.663 | 0.8718 |
| 27 | 30 | -161.4275 | 0.5848 |
| 27 | 31 | -269.7712 | 0.0481 |
| 27 | 32 | -372.3867 | 0.0021 |
| 28 | 29 | 196.2856 | 0.2579 |
| 28 | 30 | 161.5212 | 0.3986 |
| 28 | 31 | 53.1775 | 0.9968 |
| 28 | 32 | -49.4381 | 0.9986 |
| 29 | 30 | -34.7645 | 0.9999 |
| 29 | 31 | -143.1082 | 0.6738 |
| 29 | 32 | -245.7237 | 0.1045 |
| 30 | 31 | -108.3437 | 0.8491 |
| 30 | 32 | -210.9592 | 0.1710 |
| 31 | 32 | -102.6155 | 0.9122 |

Table S46: **Protocol D: White matter – sample-wise, 200 $\mu$m**

| sample1 | sample2 | diff | p.adj |
|---|---|---|---|
| 25 | 26 | 340.9691 | 0.0063 |
| 25 | 27 | 170.1757 | 0.6195 |
| 25 | 28 | -94.7193 | 0.9670 |
| 25 | 29 | 224.9304 | 0.3453 |
| 25 | 30 | 208.8066 | 0.3339 |
| 25 | 31 | 41.2462 | 0.9998 |
| 25 | 32 | -62.0061 | 0.9981 |
| 26 | 27 | -170.7934 | 0.5465 |
| 26 | 28 | -435.6885 | <0.0001 |
| 26 | 29 | -116.0387 | 0.9301 |
| 26 | 30 | -132.1625 | 0.8057 |
| 26 | 31 | -299.7229 | 0.0167 |
| 26 | 32 | -402.9753 | 0.0005 |
| 27 | 28 | -264.895 | 0.0687 |
| 27 | 29 | 54.7547 | 0.9993 |
| 27 | 30 | 38.6309 | 0.9999 |
| 27 | 31 | -128.9295 | 0.8442 |
| 27 | 32 | -232.1818 | 0.2224 |
| 28 | 29 | 319.6497 | 0.0258 |
| 28 | 30 | 303.526 | 0.0169 |
| 28 | 31 | 135.9656 | 0.7696 |
| 28 | 32 | 32.7132 | 1.0000 |
| 29 | 30 | -16.1238 | 1.0000 |
| 29 | 31 | -183.6841 | 0.5651 |
| 29 | 32 | -286.9365 | 0.0953 |
| 30 | 31 | -167.5604 | 0.5673 |
| 30 | 32 | -270.8128 | 0.0792 |
| 31 | 32 | -103.2524 | 0.9507 |

Table S47: **Protocol D: White matter – sample-wise, 500 $\mu$m**

| sample1 | sample2 | diff | p.adj |
|---|---|---|---|
| 25 | 26 | 882.1856 | <0.0001 |
| 25 | 27 | 730.5632 | <0.0001 |
| 25 | 28 | 202.7142 | 0.7373 |
| 25 | 29 | 961.2987 | <0.0001 |
| 25 | 30 | 1499.5551 | <0.0001 |
| 25 | 31 | 319.7387 | 0.1926 |
| 25 | 32 | 246.0631 | 0.5084 |
| 26 | 27 | -151.6224 | 0.8576 |
| 26 | 28 | -679.4714 | <0.0001 |
| 26 | 29 | 79.1131 | 0.9976 |
| 26 | 30 | 617.3695 | <0.0001 |
| 26 | 31 | -562.4469 | <0.0001 |
| 26 | 32 | -636.1225 | <0.0001 |
| 27 | 28 | -527.849 | 0.0001 |
| 27 | 29 | 230.7355 | 0.4969 |
| 27 | 30 | 768.992 | <0.0001 |
| 27 | 31 | -410.8244 | 0.0057 |
| 27 | 32 | -484.5001 | 0.0004 |
| 28 | 29 | 758.5845 | <0.0001 |
| 28 | 30 | 1296.8409 | <0.0001 |
| 28 | 31 | 117.0245 | 0.9604 |
| 28 | 32 | 43.3489 | 0.9999 |
| 29 | 30 | 538.2565 | 0.0003 |
| 29 | 31 | -641.56 | <0.0001 |
| 29 | 32 | -715.2356 | <0.0001 |
| 30 | 31 | -1179.8164 | <0.0001 |
| 30 | 32 | -1253.492 | <0.0001 |
| 31 | 32 | -73.6756 | 0.9974 |
